# The impact of knocking out the *Leishmania major* telomerase RNA (*Leish*TER): from altered cell proliferation to decreased parasite infectivity

**DOI:** 10.1101/2023.11.10.566567

**Authors:** Beatriz Cristina Dias de Oliveira, Mark Ewusi Shiburah, Luiz Henrique de Castro Assis, Veronica Silva Fontes, Pedro Henrique Gallo-Francisco, Selma Giorgio, Marcos Meuser Batista, Maria Nazaré Correia Soeiro, Rubem Figueiredo Sadok Menna-Barreto, Juliana Ide Aoki, Adriano Cappellazzo Coelho, Maria Isabel Nogueira Cano

## Abstract

The telomerase RNA, TER, is an intrinsic component of the telomerase ribonucleoprotein complex. It contains the telomere template sequence copied by the enzyme during telomere elongation. This unique molecule shows divergent nucleotide sequences but a more conserved secondary structure containing domains involved with telomerase assembly and biogenesis. The present work aims to characterize the biological roles played by the *Leishmania* TER component (*Leish*TER) in parasite homeostasis. We generated double knockout (*Lm*TER^-/-^) parasites, which showed a distinct growth pattern at early passages, characterized by lower density and an extended stationary phase compared to the control. Although this pattern normalized after multiple in vitro passages, ablation of *Leish*TER affected cell division and proliferation, with cells arrested at the G0/G1 phase. Progressive telomere shortening was also observed during continuous passages, along with a reduction in the expression of TERRA29. Complementation with the episomal expression of *Leish*TER did not restore telomere length to the control levels, corroborating preliminary results showing that the overexpression of TER has a dominant negative effect on parasite lifespan. *Lm*TER^-/-^ also presented a higher percentage of gamma-H2A phosphorylation, likely due to stalled replication forks since no DNA damage was observed. Also, no plasma membrane modifications were detected, but pro-survival autophagic signals were present. Intriguingly, *Lm*TER^-/-^ retained the ability to transform into metacyclic forms, although its in vitro infectivity and growth inside the host cell were compromised. Together, these results highlight the importance of TER in parasite lifespan and open a discussion about its potential as a drug target against *Leishmania*.

## Introduction

*Leishmania* spp. are unicellular eukaryotic organisms classified within the Kinetoplastida class and Trypanosomatidae family [1,2]. They present three main morphologically distinct stages of development. The procyclic promastigotes reside extracellularly in the digestive tract of insect vectors (Lutzomya or Phlebotomus species). They are highly proliferative, featuring elongated bodies and flagella that help them migrate in the insect digestive tract. At the proboscis, they transform into metacyclic promastigotes, the infective forms transmitted to mammalian hosts during the insect’s blood meal [2,3]. Metacyclic promastigotes have slender bodies with elongated flagella but are non-proliferative. After inoculating into the mammalian host, they are phagocytosed by neutrophils or macrophages. Inside the parasitophorous vacuoles, they further transform into amastigotes, characterized by rounded, highly proliferative cells with no apparent flagella [1,2,3,4]. The parasite’s ability to differentiate is primarily due to alterations in gene expression determined by environmental changes, such as pH and temperature [5].

Several *Leishmania* species cause various forms of leishmaniases^6^, characterized by diverse clinical manifestations (cutaneous, diffuse cutaneous, mucocutaneous, and visceral) with varying severity and epidemiological characteristics [6,7]. Despite their global distribution, these diseases are neglected, lacking effective treatments, human vaccines, and robust transmission control methods [8]. Understanding the molecular biology of these parasites is crucial for identifying new therapeutic targets to combat the diseases they cause. Telomeres, important for genome stability and integrity, are potential candidates for drug development [9]. They are nucleoprotein structures located at the ends of linear chromosomes in eukaryotes [10]. They avoid chromosome end degradation, recombination, and fusion and a local DNA damage response that would recognize them as DNA double-strand breaks [11]. Telomeres consist of tandemly repeated DNA sequences and exhibit a G-rich strand that extends beyond the complementary C-rich strand, forming a 3’ G-overhang [11,12]. In vertebrates and trypanosomatids, including *Leishmania*, the telomeres comprise 5’ TTAGGG 3’ repeated sequences [13,14,15].

In most eukaryotes, telomeres are maintained by telomerase, a ribonucleoprotein complex comprising two essential subunits: telomerase reverse transcriptase (TERT), which adds telomeric repeats at the 3’ G-overhangs, and Telomerase RNA (TER), a non-coding RNA that contains a template sequence used by TERT during *de novo* telomere synthesis [16,17]. TERT is highly conserved among different organisms, whereas TER is organism-specific, playing a crucial role in assembling the telomerase ribonucleoprotein complex [16,18,19,20].

TER has conserved regions, including the template sequence used by TERT and the Template Boundary Element (TBE), which prevents reverse transcription beyond the template. TBE also interacts with the TERT RNA Binding domain [21,22,23,24,25,26]. The 3’ portion of TER acts in telomerase assembly and repeat addition processivity [18,21]. In addition, it contains the pseudoknot and the Stem Terminus Element (STE) (called CR4/CR5 in humans and Helix IV in trypanosomatids) [27,28]. In some organisms, TER may contain additional domains, such as the H/ACA box in mammals and the C/D box in flagellates like trypanosomatids [13,29,30].

While the secondary structure of TER is conserved across species, the primary sequence’s size and composition vary. Despite these differences, TER has a conserved function in forming the telomerase complex and maintaining telomeres. Studies in model organisms and organisms closely related to *Leishmania* have shown that the removal or depletion of TER can lead to progressive telomere shortening, which is detrimental to cell homeostasis [25,31,32,33].

Here, we show that the knockout of both *Leish*TER alleles (*Lm*TER^-/-^) induced alterations in cell proliferation, partial G0/G1 cell cycle arrest, phosphorylation of histone γH2A, progressive telomeres shortening, and reduced expression of TERRA29. No significant morphological changes were observed in *Lm*TER^-/-^ cells, although they showed signals of an autophagic process with neither DNA fragmentation nor plasma membrane modifications. Despite presenting these molecular and cellular defects, *Lm*TER^-/-^ cells could still differentiate into metacyclic promastigotes, although impairment in infectivity compared to the parental strain was found. In addition, the overexpression of *Leish*TER also caused a dominant negative effect in *L. major*, arguing in favor of TER being important for the parasite’s homeostasis and survival.

## Results

### Generation of *Leish*TER double knockout (*Lm*TER^-/-^) was successfully achieved

Given the importance of telomerase in telomere elongation, our objective was to establish a lineage of *L. major* knockout for the TER component to investigate its impact on parasite survival. Initially, we generated an *L. major* cell lineage that episomally expresses the pTB007 plasmid (*Lm*007), encompassing the Cas9 endonuclease, T7 RNA polymerase, and the hygromycin resistance gene [34]. The *Lm*007 population was cloned by plating selection, and one clone was chosen based on Cas9 expression by Western blot (S1A Fig) and a thorough assessment of its growth pattern and cell cycle progression (S1B Fig) when compared to the wild-type parasites (*Lm*WT) (S1C Fig). Then, *Lm*007 was used as the genetic background lineage to proceed with the experiments to obtain the knockout of the two *Leish*TER alleles (*Lm*TER^-/-^) and as a control for comparative analysis with *Lm*TER^-/-^.

In the *Lm*TER^-/-^ lineages, both alleles of *Leish*TER were deleted and replaced by a donor template encoding the neomycin resistance gene, as detailed in S1E Fig.

PCR, RT-PCR, Southern blot, and Sanger sequencing were performed and confirmed the successful deletion of the two alleles of the *LeishTER* gene and its replacement by the neomycin resistance gene in the *LeishTER-*specific locus, showing that *Leish*TER knockout is not lethal to the parasites (S1D-E and S2A-D Figs, S1 Table).

### Knockout of *Leish*TER induces growth defects and arrest in the G0/G1 phase, mainly at early passages

Growth curves and proliferation assays were conducted to check if the ablation of *Leish*TER could affect the parasite development. At early passage (P5), *Lm*TER^-/-^ showed a different growth pattern than *Lm*007 in both exponential and stationary phases. At the exponential phase, the density of parasites per mL was lower than the control (Fig 1A – left panel). Despite *Lm*TER^-/-^ entering the stationary phase simultaneously with *Lm*007 (day 4), it remained longer than *Lm*007. The number of parasites was unaltered until day 12 but progressively declined (Fig 1A – left panel). Interestingly, during the continuous *in vitro* passages, *Lm*TER^-/-^ growth returned to a profile similar to *Lm*007. Growth curves of parasites at P50 (Fig 1A – right panel) showed that *Lm*TER^-/-^ promastigotes presented a similar growth profile to the parental strain *Lm*007 in exponential and stationary phases. Only on day six a significant difference at P50 was observed, and the density of control parasites decayed faster than the *Lm*TER^-/-^ (Fig 1A – right panel).

**Fig 1.**
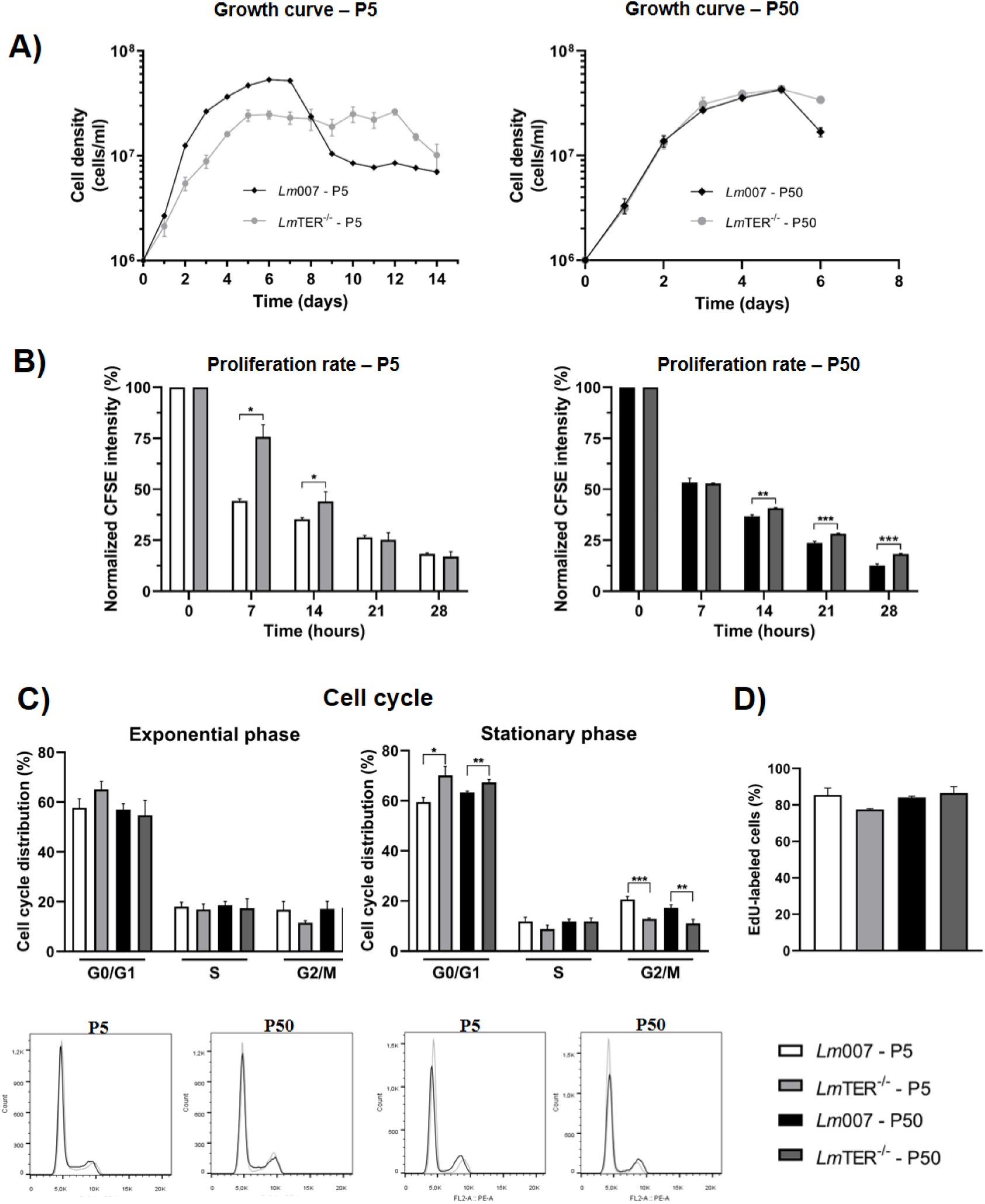
Knockout of *Leish*TER induces growth defects and arrest in the G0/G1 phase, mainly at early passages. A) Growth curves of *Lm*TER^-/-^ and *Lm*007 procyclic promastigotes at P5 (left panel) and 50 (right panel). B) CFSE labeling was used to assess the proliferation rate of *Lm*TER^-/-^ and *Lm*007 at P5 (left panel) and P50 (right panel). C) Cell cycle analysis of *Lm*TER^-/-^ and *Lm*007 in the exponential growth phase (bar graph and histograms - left panel) and stationary growth phase (bar graph and histograms - right panel) collected at P5 and P50 are depicted. D) EdU labeling was performed with *Lm*TER^-/-^ and *Lm*007 in the exponential growth phase collected at P5 and P50. All data are presented as mean ± standard deviation (S.D.) of three replicates. Statistical analysis was conducted using the student t-test with Welch’s correction when necessary. \**p* ≤ 0.05, \*\**p* ≤ 0.01, \*\*\**p* ≤ 0.001.

To explain the above results, we conducted a more sensitive proliferation assay employing Carboxyfluorescein succinimidyl ester (CFSE), which is a permeable fluorescent dye incorporated by cells and subsequently distributed among daughter cells [35], serving as a valuable tool for tracking and monitoring cell division [36]. As shown, *Lm*TER^-/-^ presented a significant reduction in CFSE fluorescence emission compared to *Lm*007, independent of which passage (P5 and P50) the parasites were analyzed. This result indicates that *Lm*TER^-/-^ had a decrease in proliferation compared to the control (Fig 1B). Briefly, *Lm*TER^-/-^ parasites from P5 were collected 7 h after adding CFSE in the culture, exhibiting a fluorescence emission of approximately 75 %, whereas *Lm*007 showed ∼45 % fluorescence under the same conditions. After 14 h, *Lm*TER^-/-^ displayed roughly 45 % fluorescence, compared to approximately 35 % in *Lm*007. This discrepancy normalized after 21-28 h (Fig 1B – left panel). At P50, *Lm*TER^-/-^ continued exhibiting higher fluorescence emission than *Lm*007 after 14 h. However, despite the statistically significant difference (*p*<0.05), the fluorescence emissions of both

*Lm*TER^-/-^ and *Lm*007 became more similar, ranging from ∼40 % (*Lm*TER^-/-^) to around 36% (*Lm*007) at 14 h, 28% (*Lm*TER^-/-^) to 23% (*Lm*007) at 21 h, and approximately 18% (*Lm*TER^-/-^) to about 12% (*Lm*007) at 28 h (Fig 1B – right panel). These results suggest that the absence of *Leish*TER impacts *L. major* promastigote cell division, with the disparity being most noticeable in *Lm*TER^-/-^ of early passages.

Further, we analyzed the cell cycle progression of non-synchronized *Lm*TER^-/-^ and *Lm*007 cultures in the exponential and stationary growth phases at P5 and P50. Independently of the passage analyzed, *Lm*TER^-/-^ and *Lm*007 in the exponential phase showed a similar cell cycle profile (Fig 1C – left panel). However, in the stationary phase, a statistically significant arrest at the G0/G1 phase was observed in *Lm*TER^-/-^ at both passages (Fig 1C – right panel). Likely, more *Lm*TER^-/-^ cells were not committed to divide compared to *Lm*007. Similar results are shown in Fig 1A-B (left panels), where *Lm*007 exhibited a higher proliferation rate than *Lm*TER^-/-^.

Cells were further incubated with EdU (5-ethynyl-2’-deoxyuridine) to monitor DNA replication. The results depicted in Fig 1D reveal that EdU incorporation is approximately 8% lower in *Lm*TER^-/-^ cells at P5 (exponential phase) compared to *Lm*007. However, this difference was not statistically significant (*p*=0.07), aligning with the outcomes of the cell cycle analysis, indicating no disparity between *Lm*TER^-/-^ and *Lm*007 in the S phase at both P5 and P50 (Fig 1C).

Altogether, the results suggest that the absence of *Leish*TER has a mild impact on promastigotes growth and cell cycle progression.

### Ablation of *Leish*TER causes progressive telomere shortening in *Lm*TER^-/-^ during continuous in vitro passages

Telomeres play a crucial role in maintaining genome stability, and it is well-established that in numerous organisms devoid of telomerase activity, telomeres tend to undergo shortening. Therefore, we initially performed a Southern blot analysis using genomic DNA from *Lm*TER^-/-^ and *Lm*007 digested with *Rsa*I and hybridized with a DIG-labeled TelC probe. It is worth reminding that in a regular telomeric Southern blot, *L. major* terminal restriction fragments appear as hybridized bands greater than 3,000 bp due to the presence of high levels of base J (β-D-glucosyl-hydroxymethyl uracil). Base J acts as an epigenetic marker, preventing DNA cleavage by restriction enzymes [37,38, 39]. Fig 2A illustrates a progressive telomere shortening of about 1.0 kb in *Lm*TER^-/-^ cells after 50 continuous in vitro passages (P5>P25>P50). The hybridization signals in *Rsa*I telomere-containing fragments decreased in size, ranging from ∼3.6 - 8.5 kb in P5 to ∼3.0 - 8.3 kb in P25 and ∼2.5 - 8.1 kb in P50. Conversely, in *Lm*007, even after 50 consecutive in vitro passages, the *Rsa*I telomere-containing fragments remained unaltered, with a size range of approximately 3.6 to 8.5 kb (Fig 2A).

**Fig 2.**
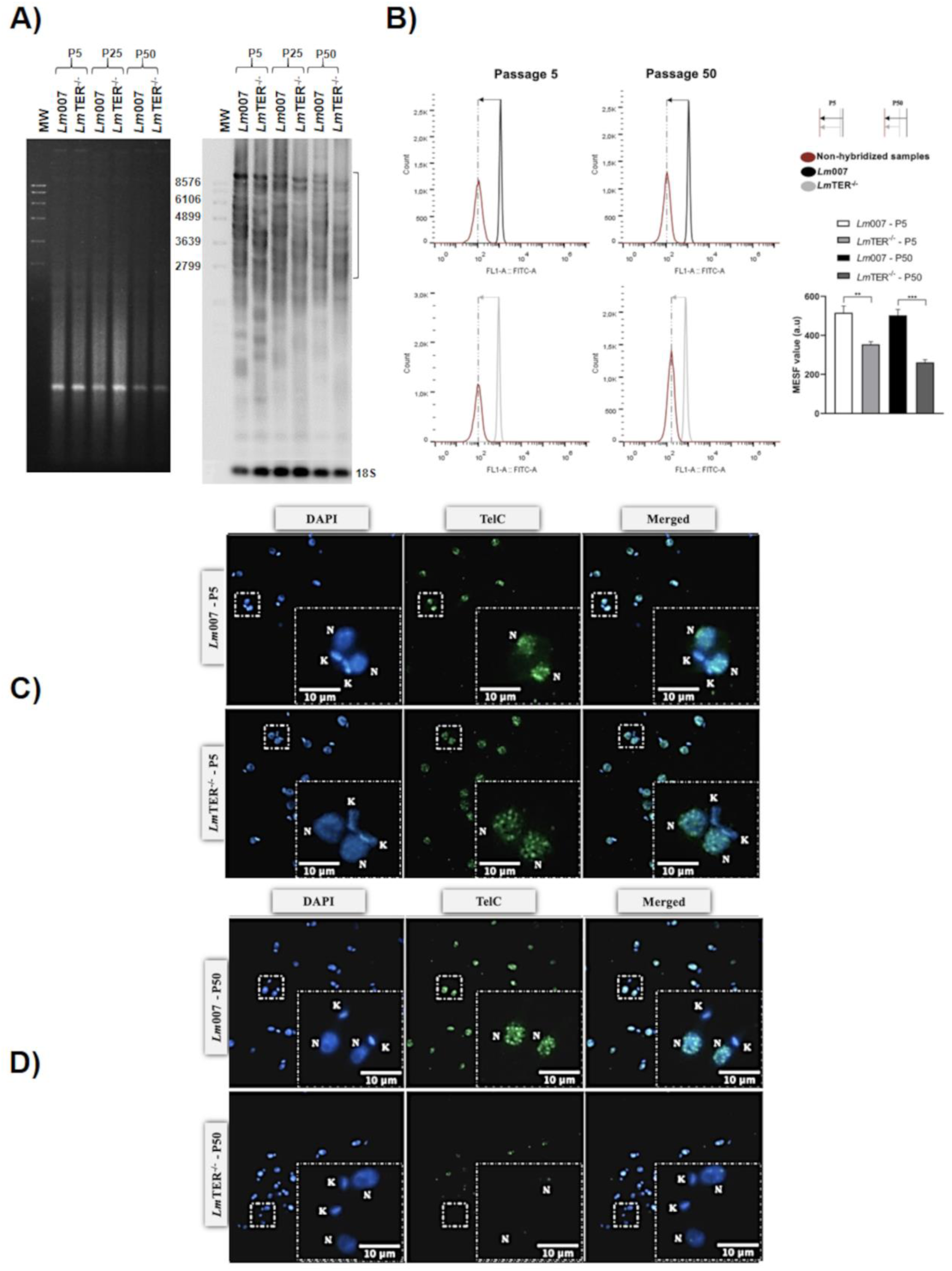
Ablation of *Leish*TER causes telomere shortening in *Lm*TER^-/-^ during continuous in vitro passages. A) Southern blot of *Rsa*I-digested genomic DNAs from *Lm*TER^-/-^ and *Lm*007 promastigotes at P5, P25, and P50, hybridized with a DIG-labeled telomeric probe (5′-TTAGGG3-3′) and a DIG-labeled 18S rRNA probe (334 bp) as the loading control. MW = DNA Molecular Weight Marker VII DIG-labeled (Roche). B) Telomeric Flow-FISH analysis of *Lm*TER^-/-^ and *Lm*007 promastigotes at P5 and P50 was done using a PNA FITC-labeled telomeric DNA oligo probe (CCCTAA)3 (Panagene). The histograms represent the mean fluorescence of non-hybridized parasites (in dark red), and parasites hybridized with the telomeric probe (*Lm*007 in black and *Lm*TER^-/-^ in gray). Vertical lines mark the difference between peaks, and horizontal lines with arrows represent telomere length differences in the populations analyzed. The amount of fluorescence among samples was calculated using MESF, and the mean ± S.D. of technical triplicate was plotted in a graph (inferior panel - left), \**p* ≤ 0.05, \*\*\**p* ≤ 0.001. C) and D) Telomeric FISH of *Lm*007 and *Lm*TER^-/-^ promastigotes at P5 and P50, respectively, hybridized with a PNA FITC-labeled telomeric DNA oligo probe (CCCTAA)3 (PANAGENE). The DNA in the nucleus (N) and kinetoplast (K) was counterstained, and the slides were mounted using VECTASHIELD® Antifade Mounting Medium with DAPI (Vector). Merged images were captured using the Nikon 80i fluorescence microscope and NIS elements software (v. Ar 3.10)—bars: 10 µm.

To perform a more accurate and quantitative analysis of the *Lm*TER^-/-^ telomere length profile, we standardized a modification of the telomeric Flow-FISH assay [40,41]. As the negative control, we used the natural fluorescence of the parasite (depicted in Fig 2B in dark red) and the fluorescence of the PNA FITC-labeled telomeric DNA oligo probe (depicted in Fig 2B in black for *Lm*007 and gray for *Lm*TER^-/-^) was used to generate the histograms. Fig 2B shows that the mean fluorescence is directly proportional to the telomere length in both *Lm*007 and *Lm*TER^-/-^ cells. The distance between the observed peaks represents the difference between telomere length in the two populations analyzed (parasites collected in P5, Fig 2B-left panel, and P50, Fig 2B – right panel). The shift towards the left in the histograms of *Lm*TER^-/-^ indicates a reduction in the emitted fluorescence, which is more pronounced in *Lm*TER^-/-^ at P50 (Fig 2B – right panel). The progressive decrease in the emitted fluorescence observed in *Lm*TER^-/-^ strongly suggests a gradual reduction in telomere length. To validate this, we calculated the molecules of equivalent soluble fluorochrome fluorescence (MESF) value and the proportional change between *Lm*007 and *Lm*TER^-/-^ (S2 Table) and plotted the results in a bar graph (Fig 2B - lower right graph). The graph highlights the gradual decrease in *Lm*TER^-/-^ telomere from P5 to P50, compared to *Lm*007.

For a more comprehensive visualization of telomere shortening, we also conducted an *in-situ* telomeric hybridization (telomeric FISH) using a PNA FITC-labeled telomeric probe (Dako). Fig 2C shows a representative analysis of about 100 cells/field. We observed that *Lm*TER^-/-^ (P5)’s telomeric signal appears more dispersed than the well-defined telomere clusters in *Lm*007. At P50 (Fig 2D), the telomeric signal in *Lm*TER^-/-^ becomes exceedingly faint, whereas in *Lm*007, the telomeric signal remains unaltered (Figs 2C-D).

Collectively, these findings undeniably indicate that the absence of *Leish*TER leads to a gradual loss of parasite telomeres with increased cell duplications (P50<P5).

### Lower expression of TERRA29 was observed in *Lm*TER^-^**^/-^** at P50

TERRA expression is known to be directly influenced by the length of the telomeres [42,43]. Hence, we investigate the potential impact of telomere shortening in the expression of TERRA from Chr29 (right arm) since it is consistently expressed in all developmental forms of the parasite [38]. RPN8 was used as the reference gene for normalizing the expression levels of TERRA29 [38].

Surprisingly, a reduction in TERRA29-R expression was observed in *Lm*TER^-/-^ cells at P50 compared to *Lm*007 (Fig 3), in disagreement with what is commonly observed in model organisms where telomere shortening induces the upregulation of TERRA expression [42,43]. In the face of this result, we intend to analyze the expression of other TERRAs to verify if the downregulation is restricted to Chr29 (right arm) or is a parasite-specific feature.

**Fig 3.**
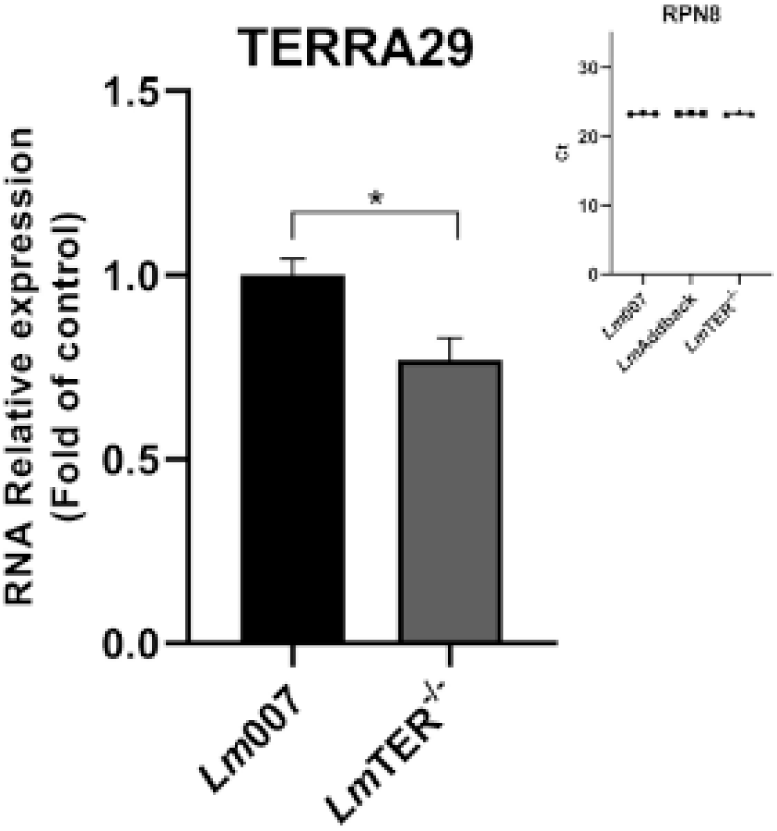
Lower expression of TERRA29 was observed in *Lm*TER^-/-^ at P50. RT-qPCR was used to estimate TERRA29 expression in *Lm*007 and *Lm*TER^-/-^ promastigotes. The expression of TERRA29 was normalized to the reference gene RPN8. *p* ≤ 0.05.

### The addback of *Leish*TER has a dominant negative effect and did not restore the telomere length to the *Lm*007 level

We conducted a complementation assay by the episomal expression of *Leish*TER in the knockout lineage, generating *Lm*Addback. PCR was performed to confirm the transfection success (Fig 4A). The restoration of *Leish*TER expression was assessed by RT-qPCR, showing high levels of *Leish*TER in *Lm*Addback, which was about 10X higher than *Lm*007 (Fig 4B).

**Fig 4.**
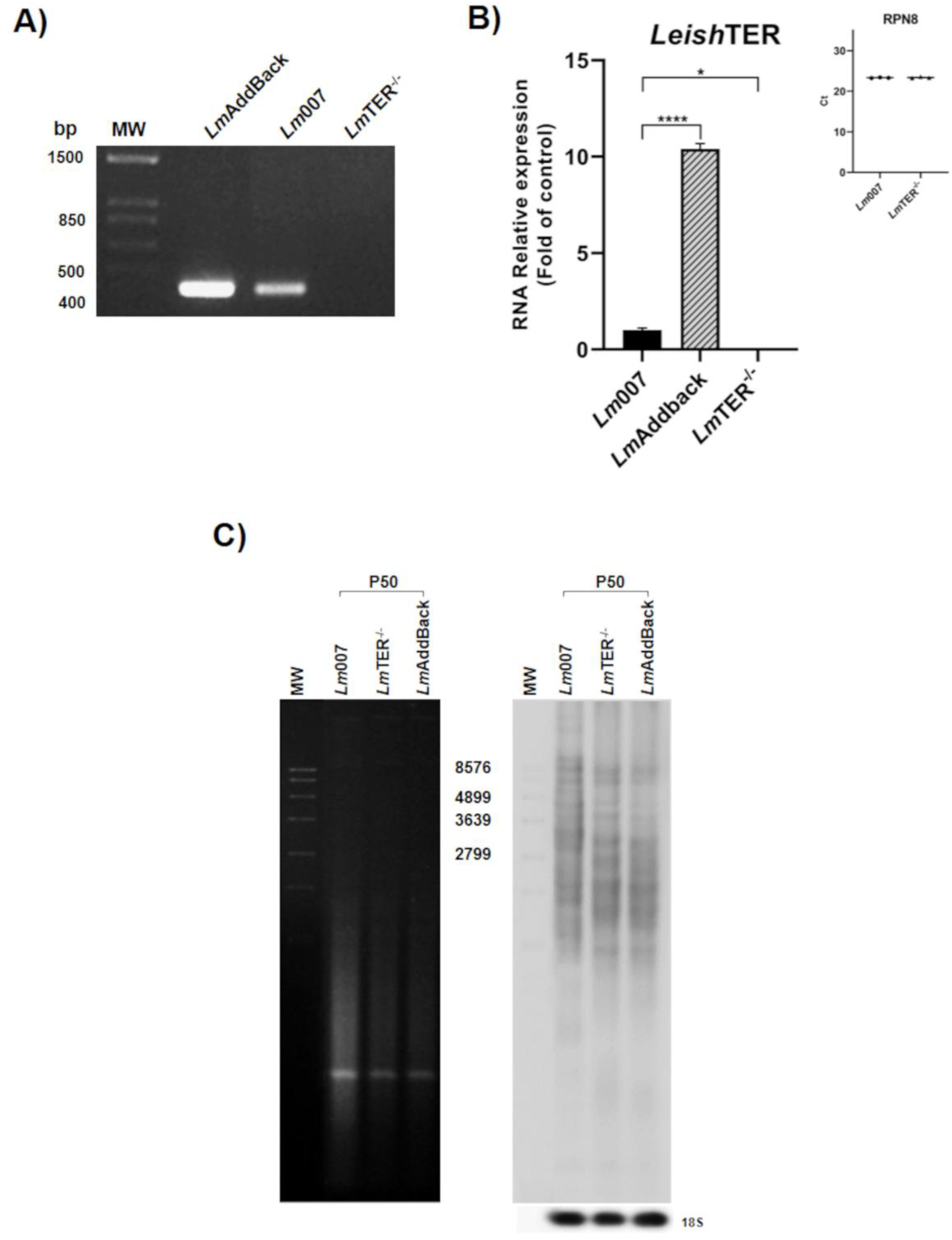
The addback of *Leish*TER has a dominant negative effect and did not restore the telomere length to the *Lm*007 level. A) PCR using genomic DNA from *Lm*TER^-/-^, *Lm*AddBack, and *Lm*007 and a specific primer set (S1 Table) to check the presence of *Leish*TER. The PCR products were separated using a 1% agarose gel stained with ethidium bromide. MW – Molecular weight 1Kb Plus DNA ladder (Invitrogen). B) To assess the expression of *Leish*TER in *Lm*007 and *Lm*AddBack, RT-qPCR was performed, with *Lm*TER^-/-^ RNA serving as the negative control. The inset displays the Ct values of RPN8, which was used as the reference gene. \*\*\**p* ≤ 0.001, \*\*\*\**p* ≤ 0.0001. C) Southern blot analysis of *Rsa*I-digested genomic DNAs extracted from *Lm*007, *Lm*TER^-/-^, and *Lm*AddBack at P50 was hybridized with a DIG-labeled telomeric probe (5′-TTAGGG3-3′) and a DIG-labeled 18S rRNA probe (334 bp) as the loading control. MW = DNA Molecular Weight Marker VII DIG-labeled (Roche).

Subsequently, parasites were grown until they reached P50 and a telomeric Southern blot was performed (Fig 4C). Curiously, the telomere length was not restored in *Lm*AddBack since the telomeric restriction profile was very similar to the *Lm*TER^-/-^ cells. Given that, we suspected that the overexpression of *Leish*TER would have a dominant negative effect on the parasite’s telomere length.

To test this hypothesis, we constructed *Lm*pXTER cells that overexpress *Leish*TER. A lineage carrying the empty pX63Neo plasmid (*Lm*pXØ) was also used as a control for the experiments. The success of the transfection was confirmed via PCR for both *Lm*pXØ and *Lm*pXTER (see S3A and B Figs). The expression of *Leish*TER was assessed through RT-qPCR, as depicted in Fig S3C, revealing that *Lm*pXTER clones expressed approximately four times more *Leish*TER than *Lm*pXØ. Upon characterization, *Lm*pXTER displayed a phenotype similar to that observed in *Lm*TER^-/-^ cells. This included alterations in the growth profiles, exhibiting lower cell density and an extended stationary phase compared to *Lm*pXØ at P5 (S3D Fig – left panel). These growth profiles were restored at P25 (S3D Fig – right panel). Regarding the cell cycle, *Lm*pXTER clones also exhibited a partial arrest at the G0/G1 phase, primarily observed at P5 (S3E Fig). Telomere shortening was observed in *Lm*pXTER, being more pronounced in clone *Lm*pXTER3 (S3F Fig; see S2 Table), in agreement with the above results.

### Phosphorylation of histone γH2A is higher in *Lm*TER^-/-^ without presenting DNA damage or apoptosis

We investigate whether telomere loss in *Lm*TER^-/-^, like in model eukaryotes, would lead to the phosphorylation of H2A [44]. Immunofluorescence assays were done using *Lm*TER^-/-^ and *Lm*007 procyclic promastigotes from P5 and P50, employing a specific anti-γH2A serum (Fig 5A). As the positive control, *Lm*007 was treated for 24 h with 10 µg/ml of phleomycin, a DNA-damage agent that induces DNA double-strand breaks [45,46,47]. Fig 5A demonstrates that γH2A was primarily detected in the nucleus of *Lm*TER^-/-^ cells from P50 (∼11%) and in the positive control (*Lm*007 treated with phleomycin) (∼47%). The bar graph in Fig 5A quantitatively represents these findings, suggesting that telomere loss in *Lm*TER^-/-^ cells can initiate phosphorylation of histone γH2A in *L. major* promastigotes just after a few population duplications, and increase after continuous in vitro passaging.

**Fig 5.**
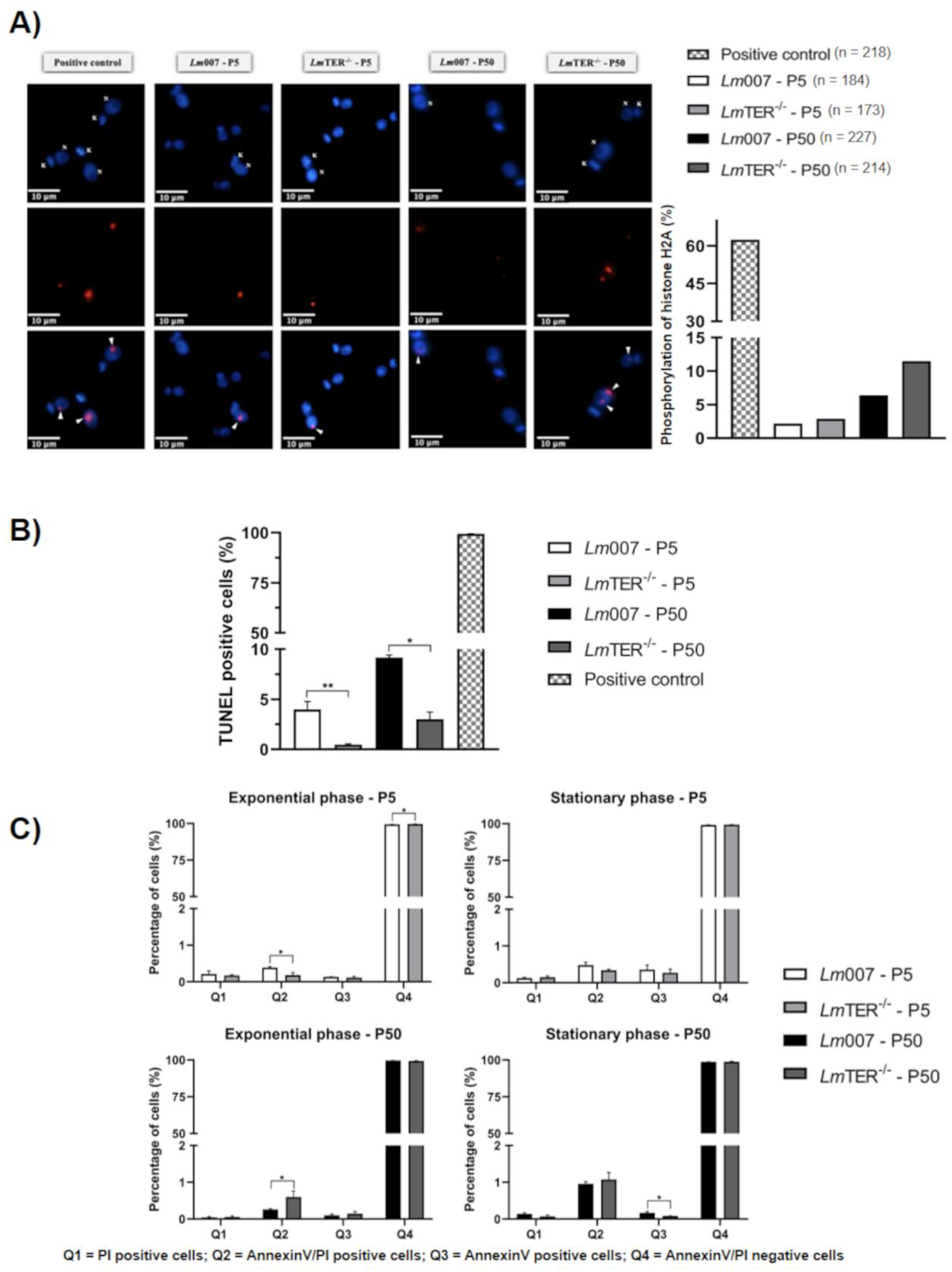
*Lm*TER^-/-^ shows higher phosphorylation of histone γH2A, although no other DNA damage or apoptosis signal was detected. A) Immunofluorescence assay was performed using *Lm*TER^-/-^ and *Lm*007 promastigotes at P5 and P50 and a specific anti-γH2A serum. As a positive control, *Lm*007 was treated with 10 µg/ml phleomycin for 24 h (hatched bar). The nucleus (N) and kinetoplast (K) were counterstained, and the slides were mounted using VECTASHIELD® Antifade Mounting Medium with DAPI. Merged images were captured using the Nikon 80i fluorescence microscope and NIS elements software (v. Ar 3.10) - scale bar: 10 μm. B) TUNEL assay was performed using *Lm*007 (white) and *Lm*TER^-/-^ (light gray) promastigotes at P5 and P50 (black and dark gray, respectively) to detect DNA fragmentation. DNaseI-treated *Lm*007 (hatched bar) served as a positive control. C) AnnexinV/PI assay was conducted on *Lm*TER^-/-^ and *Lm*007 promastigotes to assess plasma membrane integrity during the exponential and stationary phases at P5 (upper panel) and P50 (lower panel). All data are presented as mean ± SD and represent triplicate experiments. Statistical analysis was performed using the student t-test, with Welch’s correction applied when necessary. \**p* ≤ 0.05, \*\**p* ≤ 0.01.

We used the TUNEL assay to check if *Lm*TER^-/-^ cells present DNA fragmentation characteristic of cells entering apoptosis [48]. This experiment involved parasites in the exponential growth phase at P5 and P50. As shown in Fig. 5B, *Lm*007 exhibits a higher TUNEL-positive signal than *Lm*TER*^-/-^* (Fig 5B). However, the percentage of *Lm*007 cells with DNA fragmentation was low (less than 10%), suggesting a non-representative condition within the cellular population [49] (Fig 5B) and that the absence of *Leish*TER did not induce DNA damage. Besides, we checked for possible plasma membrane modifications that would signal cells entering apoptosis [50]. We assessed the integrity of the *Lm*TER*^-/-^* plasma membrane using the AnnexinV/PI assay. The results in Fig 5C indicate that approximately 1.5% of cells from both *Lm*007 and *Lm*TER^-/-^ are positive for Annexin V (Fig 5C – Q2 and Q3), with less than 0.5% showing positive PI staining, which is indicative of necrotic cells (Fig 5C – Q1). The results suggest that *Lm*TER^-/-^ does not undergo apoptosis and maintains the integrity of the plasma membrane. Therefore, it is more likely that the increased γH2A phosphorylation shown in Fig. 5A is signaling a DNA replication stress process in *Lm*TER*^-/-^* [51].

### *Lm*TER^-/-^ exhibits intracellular alterations typical of an autophagic process

To further investigate potential cellular changes induced by *Leish*TER knockout, we conducted SEM and TEM analyses on *Lm*TER*^-/-^*, as illustrated in Fig 6. Our observations reveal that, in comparison to *Lm*WT promastigotes (Fig 6A), no alteration in the *Lm*TER^-/-^ morphology could be noticed by SEM (Figs 6C and E). All samples exhibited typical features, such as elongated bodies and free flagella. Moreover, rounded and intermediate-shaped promastigotes were observed in all groups alongside actively dividing cells (Figs 6A, C, and E).

**Fig 6.**
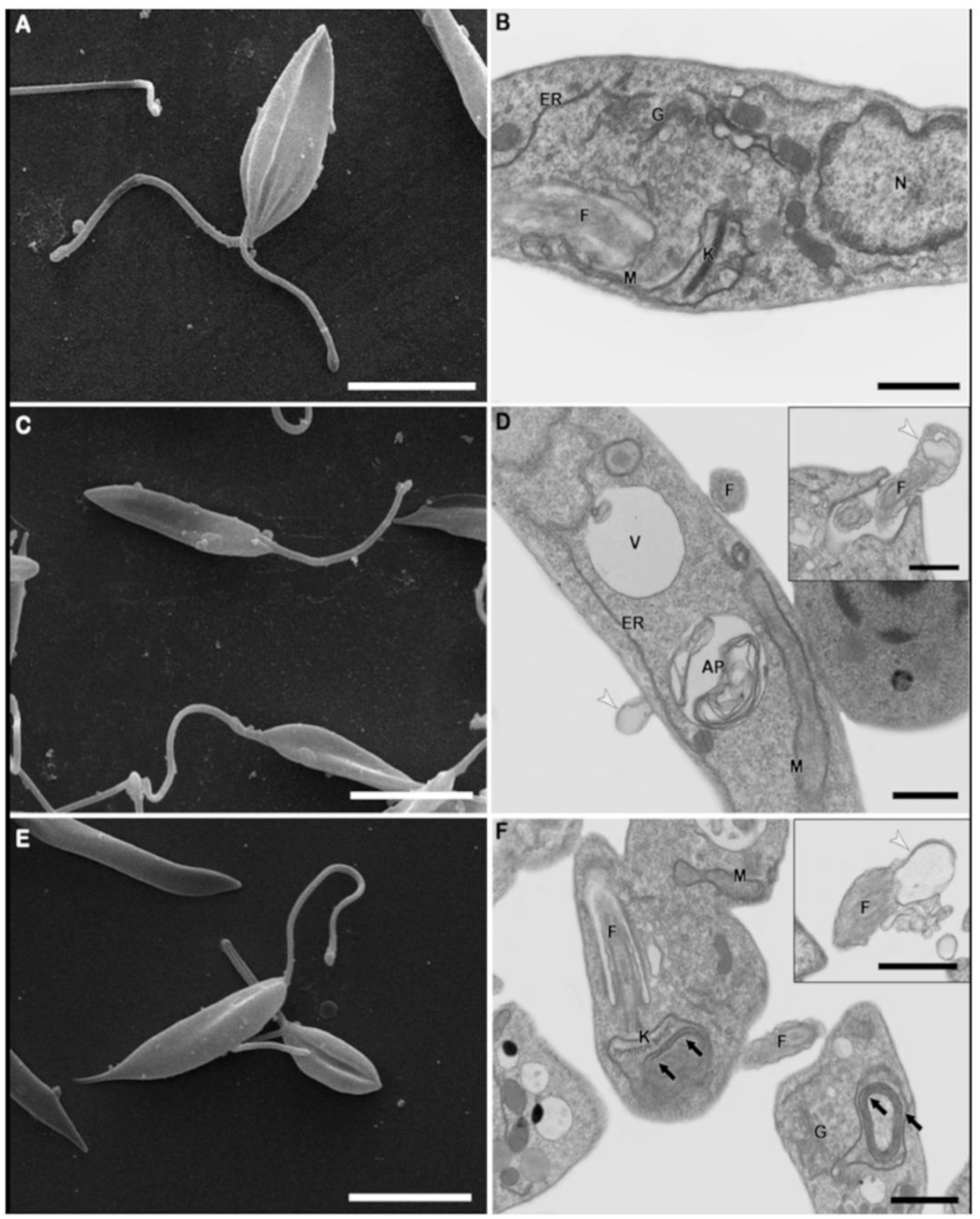
*Lm*TER^-/-^ exhibits intracellular alterations that indicate an autophagic process. SEM and TEM analyses were conducted on wild-type (A, B) and *Lm*TER^-/-^ promastigotes (C, D, E, F) at P5 (A, B, C, D) and P50 (E, F). TEM images show the nucleus (N), mitochondrion (M), kinetoplast (K), Golgi (G), endoplasmic reticulum (ER), flagellum (F), vacuole (V), autophagosomes (AP), blebs in the plasma membrane, flagella, and flagellar pocket membranes (indicated by white arrowheads) and concentric membranes within mitochondria (indicated by black arrows). Scale bars: A, C, and E: 5 µm, while B, D, and F: 1 µm.

When evaluated via TEM, the most prevalent alteration observed in *Lm*TER^-/-^ cells were intense intracellular vacuolization, formation of autophagosomes, the occurrence of plasma membrane blebs and shedding at the flagella and flagellar pocket regions besides the presence of concentric membrane structures in the cytoplasm and inside the parasite mitochondrion (Figs 6D and F). These findings strongly suggest the occurrence of an autophagic process that demands further investigation. Conversely, no parasite microtubules, nuclei, or kDNA alterations were observed in *Lm*WT and *LmTER^-/-^*promastigotes (Figs 6B, D, and F).

### *Lm*TER^-/-^ retains the ability to transform into metacyclic form, but its in vitro infectivity index is compromised

*Lm*TER^-/-^ and *Lm*007 promastigotes cultures in the stationary growth phase were subject to agglutination using peanut lectin (PNA) to select the metacyclic forms. Upon selection, a similar agglutination pattern was observed for both *Lm*007 and *Lm*TER^-/-^. The non-agglutinated metacyclic forms accounted for approximately 7-8 % of the cells (data not shown), indicating that the absence of *Leish*TER does not perturb the composition of the procyclic promastigotes plasma membrane. In contrast, parasites depleted to the TERT telomerase component present notable alterations in the procyclics plasma membrane, avoiding the parasite’s transformation into metacyclics (Shiburah & Cano, personal communication).

Subsequently, we assessed the in vitro infectivity capacity of *Lm*TER^-/-^ and *Lm*007 (P5 and P50), and *Lm*AddBack (P50) on Balb/c bone marrow-derived macrophages (BMDM) at 24 and 48 h. We chose BMDM because they show higher infection rates in vitro than other cell lines, such as THP-1 and J774.A and are better suited for observing and quantifying intracellular amastigotes [52]. Fig 7A is a representation of the in vitro infectivity assays that we performed. The graph in Fig 7B and S3 Table show that at 24 h, 30% of macrophages were infected with *Lm*007. About 18% were infected with *Lm*TER^-/-^, both at P5, whereas with parasites at P50, there was no significant difference (p >0.05) among *Lm*007 (32%), *Lm*TER^-/-^ (28%) and *Lm*Addback (36%). Furthermore, at 48 h, the percentage of macrophages infected with *Lm*007 and *Lm*TER^-/-^ at P5 increased to 69% and ∼28%, respectively. Additionally, at P50, 59% were *Lm*007-infected macrophages, and 51% and 44% were infected with *Lm*TER^-/-^ and *Lm*Addback. It is likely that the high infection rate occurred after 48 h and that independently of the parasite passage, *Lm*TER^-/-^ and *Lm*Addback infected less than the control (*Lm*007).

**Fig 7.**
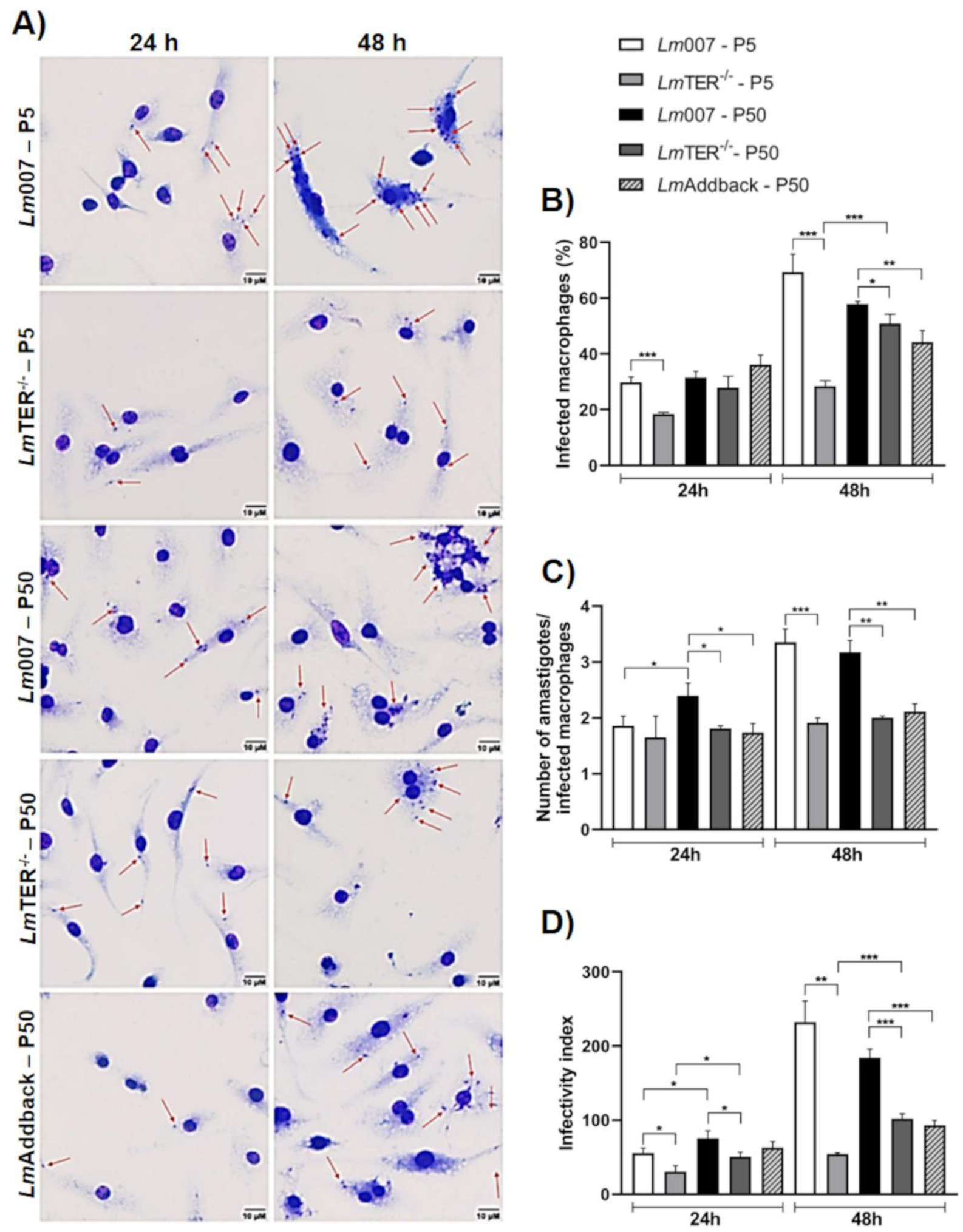
*Lm*TER^-/-^ retains the ability to transform into metacyclic form, but its in vitro infectivity index is compromised. A) Microscopic images depict bone marrow-derived macrophages (BMDM) infected for 24 and 48 h with *Lm*007 and *Lm*TER^-/-^ at P5 and P50 and *Lm*Addback at P50. Red arrows indicate *L. major* parasites internalized within the macrophages. The images were captured in representative and random fields at 40x magnification, with scale bars representing 10 µm. B) The percentage of macrophages infected for 24 and 48 h with *Lm*007 and *Lm*TER*^-/-^* at P5 and P50, along with *Lm*Addback at P50, is shown. C) Number of amastigotes per macrophage in infections with *Lm*007 and *Lm*TER^-/-^ (both at P5 and P50) and *Lm*AddBack (P50) for 24 and 48 h is shown. D) The infectivity index of *Lm*007 and *Lm*TER^-/-^ at P5 and P50 and *Lm*Addback at P50 was calculated considering the infectivity period of 24 and 48 h and are presented. All data are expressed as mean values ± S.D. and represent results from triplicate experiments. Statistical analysis was conducted using the student t-test, with Welch’s correction applied when necessary. Significance levels are indicated as \**p* ≤ 0.05, \*\**p* ≤ 0.01, \*\*\**p* ≤ 0.001.

When we estimated the number of amastigotes per infected macrophage (Fig 7C), *Lm*TER^-/-^ at P5 and P50 and the *Lm*Addback showed no difference after 24 and 48 h. Still, they all showed lower numbers compared to *Lm*007. Hence, it seems that despite the ability to infect macrophages, the proliferation of the parasites inside the host cell was impacted. Finally, the estimated infectivity index for *Lm*TER^-/-^ and *Lm*Addback at both infection periods (24 and 48 h) and passages (P5 and P50) was lower than the *Lm*007 (Fig 7D). Interestingly, there is also no statistically significant difference between *Lm*TER^-/-^ and *Lm*Addback, showing that the complementation with *Leish*TER could not restore *Lm*TER^-/-^ infectivity capacity probably because the excess of *Leish*TER exerts a dominant negative effect (see below). However, we cannot leave aside the possibility that telomere shortening (Fig 4C), which induces a decrease in parasite proliferation and cell cycle arrest (Fig 1), is implicated in a decrease in the parasite’s infectious capacity.

We also tested *Lm*TER^-/-^ infectivity capacity using the murine macrophage cell line RAW 264.7. The results post 24 h infection exhibited in S4 Fig confirm those obtained with BMDM (Fig 7), showing that *Lm*TER^-/-^ and *Lm*Addback present low infectivity capacity compared to *Lm*007.

## Discussion

Telomeres play a pivotal role in genome stability, and telomerase is essential for maintaining telomere length. In this work, we aim to understand the impact of knocking out the *Leishmania* telomerase RNA, *Leish*TER, one of the main components of the enzyme complex. Our results show the effects of *Leish*TER on parasite survival, cellular processes, and infectivity and shed light on its potential as a therapeutic target.

Our first evidence of the harm caused by the ablation of *Leish*TER in *L. major* were the effects on parasite growth, which resemble the observed in the budding yeast *Saccharomyces cerevisiae* depleted of TER and the Terc^−/−^ mouse embryonic stem cells (mESC). In yeast, TER depletion led to a decrease in growth rate and viability, whereas in mESC, the absence of Terc resulted in slow growth during extended cultures [33,53]. Our results, however, contrast with those obtained in *T. brucei,* a trypanosomatid protozoa as *Leishmania* spp.. The loss of *Tb*TER and *Tb*TERT exhibited no discernible impact on cell growth even after many population doublings (∼350) [25,54].

Furthermore, telomere shortening was detected in *Lm*TER^-/-^ during continuous in vitro passaging, aligning with the established function of telomerase in preserving telomere length. This outcome is also consistent with findings in model eukaryotes such as mice, yeasts, and *T. brucei* [25,55,56], where the depletion of TER has a detrimental impact on telomere length maintenance. Interestingly, as shown here, part of the log phase *Lm*TER^-/-^ promastigotes continued to grow in vitro until late passages (P50), suggesting that they survive even carrying short telomeres. Budding yeast depleted of TER and somatic human cells lacking telomerase activity can also survive with short telomeres [57,58]. In both cases, cells lose telomeres at every replication round until they become critically short, inducing cells to a senescence state. During senescence, cells can still avoid terminal fusions and catastrophic events of genome instability. Cells that bypass senescence, however, will face cell death or continue dividing, presenting further telomere shortening [58]. If telomeres get too short in mammals, they fuse, and cells enter crisis and eventually immortalize or transform into tumor cells. To proliferate, these cells can maintain short telomeres by reactivating telomerase or by alternative lengthening of telomeres (ALT), which usually relies on homologous recombination-mediated DNA replication [59]. In budding yeast, telomerase-depleted cells enter senescence or die after 50-100 divisions, and survivors that activate different ALT mechanisms spontaneously emerge in the cultures. Type I (∼90%) and type II (∼10%) survivors count on mechanisms of recombination/amplification to maintain the telomeric DNA tracts [57,60]. Whether *Lm*TER^-/-^ survivors use a still unknown ALT mechanism to maintain telomeres remains to be determined.

Another phenotype presented by *Lm*TER^-/-^ cells is the phosphorylation of histone γH2A, which, in trypanosomatids, differently from mammals, occurs at the Thr^130^ [61]. γ-H2A is one of the earliest markers of DNA damage, although it can also signal replication stress, including at telomeres, and cell senescence [51,62,63,64]. Since no DNA damage was detected in *Lm*TER^-/-^, we speculate that the phosphorylation of γH2A may be signaling replication stress at short telomeres, which we propose to trigger a senescence-like state in the *Lm*TER^-/-^ cells as documented by their growth impairment and G0/G1 cell cycle arrest.

Trying to revert the above phenotypes, we reintroduced *Leish*TER into *Lm*TER^-/-^ parasites (referred to as *Lm*Addback), but we did not succeed since *Lm*Addback did not restore telomere length to the control (*Lm*007) levels. Likely, the episomal expression of *Leish*TER, similar to budding yeast overexpressing TER (TLC1) [33], has a dominant negative effect inducing telomere shortening. This was further sustained by analyzing the overexpression of *Leish*TER, suggesting that an excess of *Leish*TER would disrupt either telomerase assembly and the complex stoichiometry or the episomally expressed *Leish*TER cannot acquire the proper secondary structure and, in both cases, inhibit enzyme activity.

In our study, we have also demonstrated a decreased expression of the long non-coding RNA TERRA29 in *Lm*TER^-/-^, a phenomenon that could be attributed to the shortened telomeres or the absence of *Leish*TER. While it established that short telomeres can enhance TERRA expression in mouse tumor cells and budding yeast [42,43], it’s important to mention that numerous other factors can exert influence on TERRA expression, including developmental stage, cellular stress, and telomere epigenetic modifications [43]. Therefore, further investigation is needed to determine whether the decrease in TERRA29 expression in *Leishmania* can be solely attributed to short telomeres or other factors.

Morphological transformations have also been documented in organisms lacking TER or with TER impairments. For instance, late-generation male mice deficient in TER (Terc^-/-^) exhibited testicular atrophy and depletion of male germ cells [31]. Moreover, in budding yeast, *Kluyveromyces lactis* harboring mutations in the TER, aberrant “monster cells” with varying DNA content and apparent deficiencies in cell division arose [65]. Through TEM analysis, significant intracellular changes in *Lm*TER^-/-^ parasites were observed, including initiating an autophagic process, which may indicate a mechanism for preserving cellular homeostasis [66]. However, no alterations in the DNA content and parasite plasma membrane that would indicate apoptotic events were detected.

The ability of procyclic promastigotes to transform into metacyclic promastigotes (infective forms) is key to establishing infection in the mammalian host [3,67]. Here, we show that *Lm*TER^-/-^ preserved the ability to transform into the metacyclic form, suggesting *Leish*TER is not involved in parasite differentiation. However, *Lm*TER^-/-^ exhibited a lower infectivity index than *Lm*007 across all passages and periods under scrutiny. Therefore, there appears to exist some association between reduced infectivity and telomere length. Notably, in *T. brucei,* telomerase deficient telomeres shortening increases the frequency of switch of the telomeric Variant Surface Glycoprotein-encoding genes (VSGs). VSGs are the main parasite virulent factor involved with immune evasion and are monoallelic expressed at the cell surface by a gene conversion mechanism [68]. There is no antigenic variation in *Leishmania*, but the *SCG* gene family responsible for LPG modification is mainly found near telomeres across multiple chromosomes (2, 7, 21, 25, 31, 35, and 36) [69,70]. Whether the observed telomere shortening in *Lm*TER^-/-^ affected the expression of the SCG genes should be examined.

Our results corroborate the hypothesis that the absence of *Leish*TER, besides leading to telomere shortening, may induce a senescence-like phenotype in *Leishmania* promastigotes. These findings shed light on the importance of telomerase and telomere maintenance in the survival and infectivity of *Leishmania* parasites.

Further research may explore the molecular mechanisms underlying these observations and their potential implications for leishmaniasis treatment and control.

## Materials and Methods

### Cultivation and growth curves

*Leishmania major* Friedlin strain (MHOM/IL/1980/FRIEDLIN) was used in this study. All lineages of *L. major* were cultured in M199 medium, pH 7.3 (Cultilab) supplemented with 10 % (v/v) heat-inactivated fetal bovine serum (FBS) (Cultilab) [71], and 1X penicillin/streptomycin (Life Technologies, Gibco-BRL) at 26°C. Procyclic promastigotes from passage 1 (P1) were differentiated in vitro from amastigotes extracted from mice footpad lesions [72] after inoculation into the culture medium (see above) and incubation at 26°C for 24 h. Promastigotes continuous passages represent promastigotes P1 and so on being replicated in exponential growth every three days. Metacyclic promastigotes were purified from stationary phase promastigote culture using agglutination with peanut lectin from *Arachis hypogaea* (Sigma-Aldrich), as previously described [4].

For the growth curves, the technical triplicate for each lineage was prepared with 1 x 10^6^ cells/mL. The cells were fixed in PBS containing 1 % formalin and counted in a Neubauer chamber at intervals of 24 h. The data obtained were analyzed and plotted on a graph according to the mean of the technical triplicate ± S.D., using GraphPad Prism Version 8.0.2.

### Establishment of experimental lineages

#### Obtention of *L. major* promastigotes expressing the Cas9 endonuclease and T7/RNA polymerase (*Lm*007)

*L. major* WT (*Lm*WT) procyclic promastigotes were transfected with pTB007. This episomal plasmid contains genes encoding the T7/RNA polymerase, the hygromycin phosphotransferase (hph), and the humanized *Streptococcus pyogenes* Cas9 nuclease (*hpCas9*) [34].

The transfection was performed as described by Kapler et al. (1990) [71] using the Gene Pulser Xcell Eukaryotic System (BioRad). Once cells were transfected, the presence of the pTB007 in the population was confirmed by PCR using primers that specifically amplify Cas9 and T7RNA polymerase genes (For-Cas9 + Rev-Cas9; T7RNAP-F + T7RNAP-R - S1 Table). Afterward, the parasites were plated in 1X M199/1 % agar, supplemented with 10 % (v/v) FBS and 30 μg/mL hygromycin to select clones. Clones of *L. major* strain expressing Cas9 and T7/RNAp (*Lm*007) were inoculated in 1X M199 liquid medium supplemented with 10 % (v/v) FBS and 30 μg/mL hygromycin B (Invitrogen).

#### Generation of *Leish*TER knockout (*Lm*TER^-/-^) lineage

A donor DNA cassette that confers resistance to neomycin/G418 and two sgRNAs were used to obtain *Lm*TER^-/-^ cells. For the in vivo transcription of guide RNAs (one for the 5’ end and the other for the 3’ end of *Leish*TER), sgRNA templates were generated by PCR following the protocol described in Beneke et al. (2017) [34] using both sets of primers: TracrRNA (G00) – scaffold + TER seed-5’ and TracrRNA (G00) – scaffold + TER seed-3’ and (see S1 Table for the sequences of the primers). The sgRNA sequences were chosen using the online tool Eukaryotic Pathogen CRISPR guide RNA/DNA Design Tool, EuPaGDT (http://grna.ctegd.uga.edu) using the default parameters described in Beneke et al. (2017) [34]. The donor DNA was also obtained by PCR using the pTNeo plasmid as a template [34]. Transfections were performed as described earlier [71]. LmTER-/- clones were selected using solid M199 medium, as described in the previous item. *Lm*TER^-/-^ cells were maintained with 40 µg/mL G418 (Sigma-Aldrich).

#### Selection of *Lm*TER^-/-^ clones using PCR, RT-PCR, and automated sequence

A set of primers that hybridizes at the 5’ upstream and 3’ downstream sequences of the *Leish*TER gene (5’TER F + 3’TER R - S1 Table; S1D and E Figs) were used to confirm the generation of the knockout line. The above primers set would generate an amplicon of 2,432 bp *Lm*007 used as control and an amplicon of 2,059 bp if the donor DNA cassette replaced the *Leish*TER gene in the correct locus.

Total RNA obtained from the *Lm*TER^-/-^ and from the control (*Lm*007) and a set of primers that hybridizes in the internal portion of *Leish*TER gene (TERmF and TER RT R - S1 Table; S1D Fig) were used to obtain double-stranded cDNA using SuperScript™ III One-Step RT-PCR System with Platinum™ Taq DNA Polymerase (Invitrogen), according to manufacturer’s instructions.

Sequencing reactions were performed using primers that target the regions upstream and downstream of the *Leish*TER locus and internal regions of the donor DNA (S1E Fig and S1 Table). Fragments of interest were generated by PCR with further purification using the PureLink™ PCR Purification Kit (Invitrogen), and then automated Sanger sequencing was performed at IBTEC (UNESP – Botucatu). Blastn, Clustal Omega online tool (https://www.ebi.ac.uk/Tools/msa/clustalo/) and SnapGene 6.0.2. were used for in silico sequence analysis.

#### Complementation assay: addback of *Leish*TER in *Lm*TER^-/-^ cells

*Lm*TER^-/-^ cells were transfected with the plasmid pSP72-a-bla-a-LmjTER synthesized by GenOne using the pSP72-a-bla-a plasmid as the backbone. First, the transfection was performed, and 20 µg/mL of blasticidin (Invivogen) was added to the culture to select the transfectants. Then, PCR (Primers: TERmF + TER RT R – S1 Table) and RT-qPCR (Primers: TER RT F + TER RT R – S1 Table) were performed to confirm the presence of the plasmid and expression of *Leish*TER, respectively. *Lm*AddBack was maintained in 40 µg/mL G418 and 20 µg/mL Blasticidin (Invivogen).

#### Generation of an *L. major* lineage overexpressing *Leish*TER

*L. major* at the exponential growth phase were transfected with 50 μg of *Leish*TER-pX63Neo plasmid DNA (Vassilievitch & Morea, *unpublished*), which contains the *Leish*TER gene and the gene that confers resistance to neomycin. In addition, *Lm*WT was also transfected with 50 µg of empty pX63Neo plasmid [73] as a control. The pX63Neo plasmid is maintained as an episome in *Leishmania* with 40 μg/mL of G418. It is specifically used for expressing *Leishmania* genes since it signals trans-splicing and polyadenylation [73]. The selection of transfectants containing *Leish*TER-pX63Neo and pX63Neo plasmids (named *Lm*pXTER and *Lm*pXØ, respectively) was performed using 40 μg/ml G418 as the selection antibiotic. The genotype of the clones was determined by PCR, using specific primers (Px63 Neo Fw and Px63 Neo Rv - S1 Table), and the overexpression of *Leish*TER was confirmed by RT-qPCR (Primers: TER RT F and TER RT R - S1 Table).

### DNA extraction and Southern blot

Genomic DNA was obtained from 2 x 10^8^ cells of *L. major* using DNeasy Blood and Tissue kit (Qiagen) according to the manufacturer’s instructions and phenol:chloroform extraction, according to Medina-Acosta and Cross (1993) [74] with some modifications as follows: 2 x 10^8^ parasites were washed twice with 1X PBS and resuspended in TELT buffer (50 mM Tris-HC1, pH 8.0; 62.5 mM EDTA, pH 9.0; 2.5 M LiCl and 4% (v/v) Triton X-100) containing 10 μg/μl proteinase K and incubated at 56 °C for 10 min. Subsequently, 1 volume of phenol:chloroform:isoamyl alcohol (25:24:1) was added, and the tube was inverted for 5 min, followed by centrifugation at 16,000 g for 10 min. The supernatant was transferred to a new tube, and 1 ml of 100% cold ethanol was added. The tube was inverted a few times until the DNA was precipitated and centrifuged at 16,000 g for 5 min. The pellet of DNA was washed once with 70% cold ethanol, centrifuged, and air-dried. After that, the samples were resuspended in 100 µl of TE (10 mM Tris-HCl, pH 8.0/11 mM EDTA, pH 8.0) containing 50 μg/μl RNase A (Invitrogen) and incubated at 37 °C. DNA concentration and purity were evaluated by calculating the A260/280nm ratio using a spectrophotometer (Epoch, BioTek). DNA integrity was verified on 1% agarose gel.

For Southern blot, 1 μg of genomic DNA was digested with 5U *Rsa*I (Roche) at 37 °C overnight to release the telomeric fragments [75]. The subsequent steps of the Southern blot were done using standard procedures [75, 76] with minor modifications as described in Morea et al. (2021) [38]. For the confirmation of *Leish*TER ablation in *Lm*TER^-/-^, DIG-labeled TER and alpha-tubulin probes were generated using the PCR DIG Probe Synthesis Kit (Roche) and used in the hybridization step (S1 Table, S1D Fig).

Southern blot membranes were hybridized using a DIG-labeled TELC probe (S1 Table) or a DIG-labeled 18S rRNA probe (S1 Table) as the loading control. The Telomere Restriction Fragments (TRF) location on the blots was compared to the DNA molecular weight marker VII DIG-labeled (Roche) to obtain the average of TRF.

### Total protein extraction and Western blotting

Protein extracts were obtained according to de Oliveira et al. (2021) [77] using 5×10^8^ cells harvested at 2,300 g for 5 min and washed twice with 1X PBS. The pellet was resuspended in supplemented buffer A (20 mM Tris-HCl pH 7.5; 1 mM EGTA pH 8.0; 1 mM EDTA pH 8.0; 15 mM NaCl; 1mM spermidine; 0.3 mM spermine; 1mM DTT) and incubated in liquid nitrogen for 1 minute followed by incubation on ice until defrost. A solution containing 0.5 % of Nonidet P-40 (Calbiochem) and 1X protease cocktail inhibitor (Sigma-Aldrich) was added to the samples, followed by 30 min incubation on ice. The samples were then centrifuged at 16,000 g for 20 min at 4 °C; subsequently, the supernatant was transferred to a new tube, and the total protein in each extract was estimated by reading the OD280_nm_ using the spectrophotometer (Epoch, BioTek). For the Western blot, 200 μg of total protein extracts were fractionated in a 10 % SDS-PAGE followed by a wet transfer to a PVDF membrane (GE Healthcare) [78]. The membranes were probed with primary antibodies, Anti-Flag (dilution 1:5,000 – Sigma-Aldrich) and Anti-*Lm*GAPDH (dilution 1:2,000 - GenScript), followed by incubation with Anti-rabbit IgG (dilution 1:50,000 - BioRad) as the secondary antibody. According to the manufacturer’s instructions, immunoreactive bands were revealed by using ECL Prime Western Blotting Detection Reagent (GE Healthcare).

### Total RNA isolation and RT-qPCR

Total RNA from 3 x 10^8^ cells at the exponential growth phase was isolated using TRIzol reagent (Invitrogen). RNase-free DNase I treatment (Thermo Scientific™) was conducted, followed by inactivation through incubation with 1 µl of 25 mM EDTA at 65 °C for 10 min, according to the manufacturer’s instructions. RNA concentration and purity were evaluated by an A_260nm_ (spectrophotometer Epoch, BioTek). The absence of genomic DNA in the RNA samples was confirmed by PCR using primers for alpha-tubulin (Primers: Alfa Fw1 + Alfa Rv1 - S1 Table) and positive control (*Lm*WT genomic DNA).

First-strand cDNA synthesis was performed using the iSCRIPT cDNA synthesis kit (BioRad) according to the manufacturer’s instructions. The RT-qPCR assays were performed in technical triplicate using cDNA, iTaq Universal SYBR® Green Supermix (BioRad), and 0.5 μM of each primer specific for *Leish*TER and the subtelomeric region of chromosome 29 - right arm (S1 Table). The gene encoding the 19S proteasome non-ATPase subunit 8 (RPN8) (*Lmj*F.32.0390) was used as a control [38, 79] to normalize gene expression levels of the targets (*Leish*TER and TERRA29). RT-qPCRs were performed in 10 μL reactions, and cycle threshold (Ct) values were obtained in StepOne™ Real-Time PCR System (Applied Biosystems) using default settings. The relative gene expression data were analyzed using the 2^−ΔΔCT^ method as described [80]. The data were expressed as means of fold change over control (relative expression), where the control data was normalized to an arbitrary value of 1. Statistical analyses were performed using GraphPad Prism version 8.0.2.

### Flow cytometry protocols

#### DNA content analysis

1.5 x 10^7^ cells at the exponential and stationary growth phase were harvested at 900 g for 5 min, washed twice with 1X PBS, fixed and permeabilized with 90% ice-cold methanol, followed by incubation at −20 °C for 20 min as described in Barak et al. (2005) [81] with some modifications [82]. Then, cells were washed once with 1X PBS, resuspended in staining solution (1X PBS containing 100 μg/mL of RNase-A (Invitrogen) and 10 μg/mL of propidium iodide (Thermo Scientific™), and incubated for 40 min at 37 °C. Subsequently, the parasites were analyzed using flow cytometry (Acuri C6 Plus – BD Bioscience), and the DNA content analysis was performed according to de Oliveira et al. (2022) [82]. Technical triplicates were used, and 20,000 events were analyzed for each sample.

#### Cell proliferation tracking using carboxyfluorescein succinimidyl ester (CFSE)

1 x 10^7^ cells at the exponential growth phase were labeled with 8 µM CFSE (Invitrogen) following the manufacturer’s instruction. The first 20 min after labeling was considered time zero, with parasites emitting 100 % CFSE fluorescence. Then, parasites were left in culture and collected every 7 h up to 28 h, and CFSE fluorescence emission was monitored using flow cytometry [83]. About 7,000 events of each sample were analyzed. The mean fluorescent intensity (MFI) was estimated and normalized, and the mean of technical triplicate ± standard deviation was plotted in GraphPad Prism Version 8.0.2.

#### Telomeric Flow-FISH

Telomeric flow-FISH was performed using a modification of the Baerlocher et al. (2006) [40] method as described by da Silva et al. (2017) [41] without the human leukocytes: 1.5 x 10^7^ cells were centrifuged at 900 g for 5 min at 4 °C and washed twice with 1X PBS, fixed and permeabilized with 90% ice-cold methanol, followed by incubation at −20 °C for 20 min. Then, cells were washed twice with 1X PBS (5,000 g for 5 min at 4 °C), and the pellet was gently resuspended in hybridization buffer (70 % formamide, 20 mM Tris-HCl pH 7, and 1 % BSA) containing a PNA FITC-labeled telomeric DNA oligo probe (CCCTAA)_3_ (Panagene). The pellet was gently resuspended only in the hybridization buffer for the non-hybridized controls. Samples were protected from light and incubated at 85 °C for 15 min, followed by overnight (∼16h) incubation at room temperature. Parasites were washed in 1X Wash Solution (Dako) and incubated for 10 min at 40 °C, then centrifuged at 5,000 g for 5 min. The supernatant was discarded, and this step was repeated, followed by an additional wash with 1X PBS and centrifugation at 5,000 g for 5 min. The pellet was resuspended in Staining solution (1X PBS containing 100 μg/mL of RNase-A and 10 μg/mL of propidium iodide) and incubated for 30 min at 37 °C and then for 30 min at 4 °C. Samples were subjected to flow cytometry analysis using Accuri C6 Plus (BD Bioscience). Quantum™ FITC-5 MESF beads (BioRad) were used according to the manufacturer’s protocol to estimate the length of telomeres quantitatively. Technical triplicates of hybridized and non-hybridized telomeres were used. For each sample, approximately 20,000 events were analyzed.

#### Verification of DNA fragmentation and plasma membrane modifications by TUNEL and Annexin-V, respectively

For the TUNEL assay, 1 x 10^7^ cells at P5 and P50 at the exponential growth phase were used following the manufacturer’s instruction for cells in suspension. As a positive control, cells were treated with 10 units/mL Deoxyribonuclease I from bovine pancreas (Sigma-Aldrich) for 10 min. As the negative control, cells were prepared without incubation with rTdTenzyme, following the manufacturer’s instructions (DeadEnd™ Fluorometric TUNEL System – Promega).

The Annexin-V assay followed the manufacturer’s protocol (BioRad) using 1 x 10^6^ cells. In addition, three technical replicates of *Lm*007 and *Lm*TER^-/-^ promastigotes in the exponential and stationary growth phase at P5 and P50 were used. A sample not labeled with AnnexinV-FITC and PI was used as the negative control. As a positive control, cells were treated with 3 % formaldehyde for 30 min on ice before incubation with AnnexinV:FITC. Approximately 10,000 events were collected to be analyzed in a flow cytometer.

#### Assessment of DNA replication by EdU labeling

Following the standard procedures as described by da Silva et al. (2017) [84], cells at exponential growth phase were incubated with 200 μM of the thymidine analog EdU (5-ethynyl-2′-deoxyuridine - Invitrogen) at 26 °C for 10.2 h (estimated doubling time of *L. major*) [85] followed by centrifugation at 900 g at 4°C for 5 min. The supernatant was discarded, and the pellet was washed twice with 1X PBS. The supernatant was discarded carefully and resuspended in cold 1X PBS, followed by the addition of ice-cold methanol (final concentration of 90%). After mixing by pipetting, the samples were incubated at −20°C for 20 min for fixation. Afterward, samples were vortexed and washed once with 1X PBS. 100 μL of cycloaddition reaction (3 mM CuSO4, 100 mM Ascorbic Acid, and 5 mM Alexa Fluor azide 488) was added in each sample and incubated for 1 h protected from light. Samples were washed once with 1X PBS, and the pellet was resuspended in 500 μL of 1X PBS. Approximately 20,000 events were collected from each sample by flow cytometry.

#### Sample collection and analysis using Flow cytometry

The sample collections were done using the Accuri C6 Plus flow cytometer (BD Bioscience), and the data analysis was done using both FlowJo v.10.6.5 software and GraphPad Prism Version 8.0.2. Two standard graphs were generated at FlowJo for all experiments: FSC-A x SSC-A and FSC-A and FSC-H to select the population of interest and exclude doublets and debris. In addition, due to the particularity of each experiment, additional graph was generated to analyze the fluorescence emission related to each experiment as follow: for DNA content analysis: FL2-A x FL2-H and FL2-A x histogram; for TUNEL: SSC-A x SSC-H, FL2-A x FL2-H and FL1-A x histogram, Flow-FISH: FL2-A x FL2-H and FL1-A x histogram; for CFSE and EdU: FL1 x histogram; and Annexin V: FL2-A x FL1-A.

### Telomeric FISH

For these assays, we used a protocol described before [77]. 1 x 10^7^ cells from P5 and P50 were fixed with 4% paraformaldehyde for 5 min on ice, followed by several washes in 1X PBS. Then, the fixed cells were attached to glass slides coated with 0.1% poly-L-lysine (Sigma-Aldrich). Subsequently, cells were dehydrated through an ice-cold ethanol series (70 %, 85 %, and 100 %) for 2 min each and air dried. Cells were incubated for 5 min at 85 °C and then 30 min at room temperature, protected from light, with 10 μM of PNA FITC-labeled telomeric DNA oligo probe (CCCTAA)3 (Panagene). After the hybridization step, cells were washed once with Rinse solution (Dako) at room temperature for 10 seconds, followed by another washing using 1X Wash solution (Dako) at 65°C for 5 min. Afterward, a second dehydration step was followed, as described earlier. The DNA in the nucleus and kinetoplast was counterstained, and the slides were mounted using VECTASHIELD® Antifade Mounting Medium with DAPI (Vector). Approximately 100 cells/field were observed in each experiment. The slides were analyzed under a Nikon 80i fluorescence microscope, and the images were captured using the NIS elements program (Version Ar 3.10).

### Detection of histone **γ**H2A phosphorylation

The immunofluorescence assay was performed as described before [41]: 1 x 10^7^ cells were harvested at 900 g for 5 min at 4°C, washed with 1X PBS, fixed in 4% paraformaldehyde, washed with 1X PBS, and then placed on a glass slide previously coated with 0.1 mg/ml of poly-L-lysine (Sigma-Aldrich). Cells were subsequently permeabilized with 0.1% Triton X-100 for 5 min, washed several times, and then incubated overnight at 4°C with the primary antibody anti-γH2A (a gift from Dr. Richard McCulloch - University of Glasgow) (dilution 1:500). Slides were washed with 1X PBS and incubated with anti-rabbit IgG AlexaFluor 594 (Invitrogen) for 2 h protected from light followed by three washes using 1X PBS. The DNA in the nucleus and kinetoplast was counterstained, and the slides were mounted using VECTASHIELD® Antifade Mounting Medium with DAPI (Vector). *Lm*007 treated with 10 µg/ml of phleomycin (Invivogen) for 24 h was used as a positive control [45]. The slides were analyzed under a Nikon 80i fluorescence microscope, and the images were captured using the NIS elements program (Version Ar 3.10).

### Transmission (TEM) and Scanning Electron Microscopy (SEM) analysis

*L. major* procyclic promastigotes (*Lm*WT and *Lm*TER^-/-^) at P5 and P50 were washed twice with 1X phosphate-buffered saline (PBS) and fixed at 4 °C for 40 min with 2.5 % glutaraldehyde (GA - Sigma-Aldrich) diluted in 0.1 M 2.5 % glutaraldehyde in 0.1 M Na-cacodylate buffer (pH 7.2). The parasites were prepared for TEM analysis as described previously [86]. Ultrathin sections were stained with uranyl acetate and lead citrate and examined under a Jeol 1200 EX transmission electron microscope (JEOL USA Inc.) at Centro Nacional de Biologia Estrutural e Bioimagem (CENABIO, Rio de Janeiro).

For SEM analysis, the GA-fixed parasites were submitted to the procedures described by Rebello et al. (2013) [87] and examined with a Jeol JSM6390LV scanning electron microscope (JEOL USA Inc.) at Instituto Oswaldo Cruz Platform [87].

### In vitro macrophage infection

#### In vitro infection using bone marrow-derived macrophages

Bone marrow-derived macrophages (BMDMs) were obtained from the femurs and tibias of BALB/c mice, which were differentiated *in vitro* as previously described [88].

A total of 3×10^5^ cells were seeded onto sterile coverslips (Olen) in 24-well plates (Nest) and incubated overnight at 37 °C and 5 % CO_2_ in RPMI medium as previously described [89]. Non-adherent cells were removed, and the infection was performed with *Lm*007, *Lm*TER^-/-^ (P5 and P50), and *Lm*AddBack (P50) parasites in the stationary growth phase (MOI 10:1) for 3 h at 34°C and 5% CO2. After this period, non-internalized parasites were removed by washing with warmed 1X PBS, and then infections were maintained for 24 or 48 h. Following the infection, the infected macrophages were fixed with methanol and stained using a Panoptic kit (Laborclin). The experiments were carried out in triplicate, and the observations were made at a magnification 40X of the optical light microscope (Leica DME). In addition, photomicrographs were taken using the EVOS XL Core Imaging System (ThermoFisher) at 40X magnification.

#### In vitro infection using RAW 264.7 murine macrophages

A suspension of 2 x 10^5^ macrophages/mL of RAW 264.7 murine macrophages ATCC TIB-71™ (Rio de Janeiro Cell Bank – BCRJ) was cultured on 24-well plates containing 13 mm diameter glass coverslips in complete media (RPMI-1640 supplemented with 10 % FBS, 2 mM L-glutamine and 10 mM HEPES) and incubated overnight at 37 °C, 5 % CO_2_, 21 % of balanced O_2_ and N_2_ for macrophage step adhesion. Afterward, the macrophage culture was infected with *Lm*007 and *Lm*TER^-/-^ (P5 and P50) and *Lm*Addback (P50) transgenic lines in the stationary growth phase, as described previously [90]. We used a ratio of MOI 10:1 for infections. After the incubation period (24 h), the coverslips were fixed with methanol and stained with Giemsa to visualize the macrophages and the internalized parasites. The experiments were carried out in triplicate. The observations were made at a magnification of 1,000x of the optical light microscope (Zeiss Primo Star, Oberkochen, GER). In addition, photomicrographs were taken on a camera (Axiocam ERc5s, Axiovision software 4.8, Zeiss) attached to the microscope.

#### Infection analysis

The percentage of infection was calculated by dividing the number of infected macrophages by the total number of macrophages. The average number of amastigotes per infected macrophage was determined by dividing the total number of amastigotes by the total number of infected macrophages. The infectivity index was calculated by multiplying the percentage of infected macrophages by the mean number of intracellular amastigotes per infected macrophage [91].

### Statistical analysis

The mean and standard deviation (S.D.) were calculated from a technical triplicate of each sample to be analyzed. A two-tailed non-parametric unpaired Student’s *t*-test was performed when it was established that the variances among the groups were homogenous following an examination for equal variance. In cases where a significant variance difference was detected, Welch’s t-test was employed. A probability level of 0.05 was chosen for statistical significance, where \**p* ≤ 0.05, \*\**p* ≤ 0.01, \*\*\**p* ≤ 0.001, and \*\*\*\**p* ≤ 0.0001. GraphPad Prism Version 8.0.2 was used to perform all the statistical analyses and to generate the graph.

## Data availability

This paper does not report any original code.

All data reported in this paper can be found at doi:10.5281/zenodo.10035539, along with the respective statistical treatments and results obtained.

Any additional information required to reanalyze the data reported in this paper is available from the lead contact upon request.

## Ethics statement

Animal procedures were approved by the Ethics Committee for Animal Experimentation of the Instituto de Biologia, Universidade Estadual de Campinas (UNICAMP) (protocol: 5719–1/2021) for experiments using mice.

## Acknowledgments

The authors want to acknowledge Dr. Elton José R. Vasconcelos for the critical reading of the manuscript. Dr. Marcelo Santos da Silva, Dr. Débora Andrade da Silva, and Miss Stephany Cacete de Paiva for the comments and directions of some experiments and methodology.

## Author contributions

**Conceptualization:** M.I.N.C., **Data curation:** B.C.D.O., M.I.N.C., **Formal analysis:** B.C.D.O., P.H.G.F., **Funding acquisition:** M.I.N.C., **Investigation:** B.C.D.O., M.E.S., L.H.C.A., V.S.F., P.H.G.F., M.M.B., J.I.A., **Methodology:** B.C.D.O., M.E.S., L.H.C.A., P.H.G.F., M.M.B. J.I.A., **Project administration:** M.I.N.C., **Resources:** M.I.N.C., S.G., M.N.C.S., R.F.S.M.B., A.C.C., **Supervision:** M.I.N.C., **Validation:** B.C.D.O., **Visualization:** B.C.D.O., S.G., M.N.C.S., **Writing – Original draft:** B.C.D.O., P.H.G.F., **Writing – Review & Editing:** B.C.D.O., M.E.S., L.H.C.A., P.H.G.F., S.G., M.N.C.S., J.I.A., A.C.C., M.I.N.C.

