## Supplemental file for "The impact of knocking out the *Leishmania major* telomerase RNA (*Leish*TER): from altered cell proliferation to decreased parasite infectivity"

**Supplementary material**

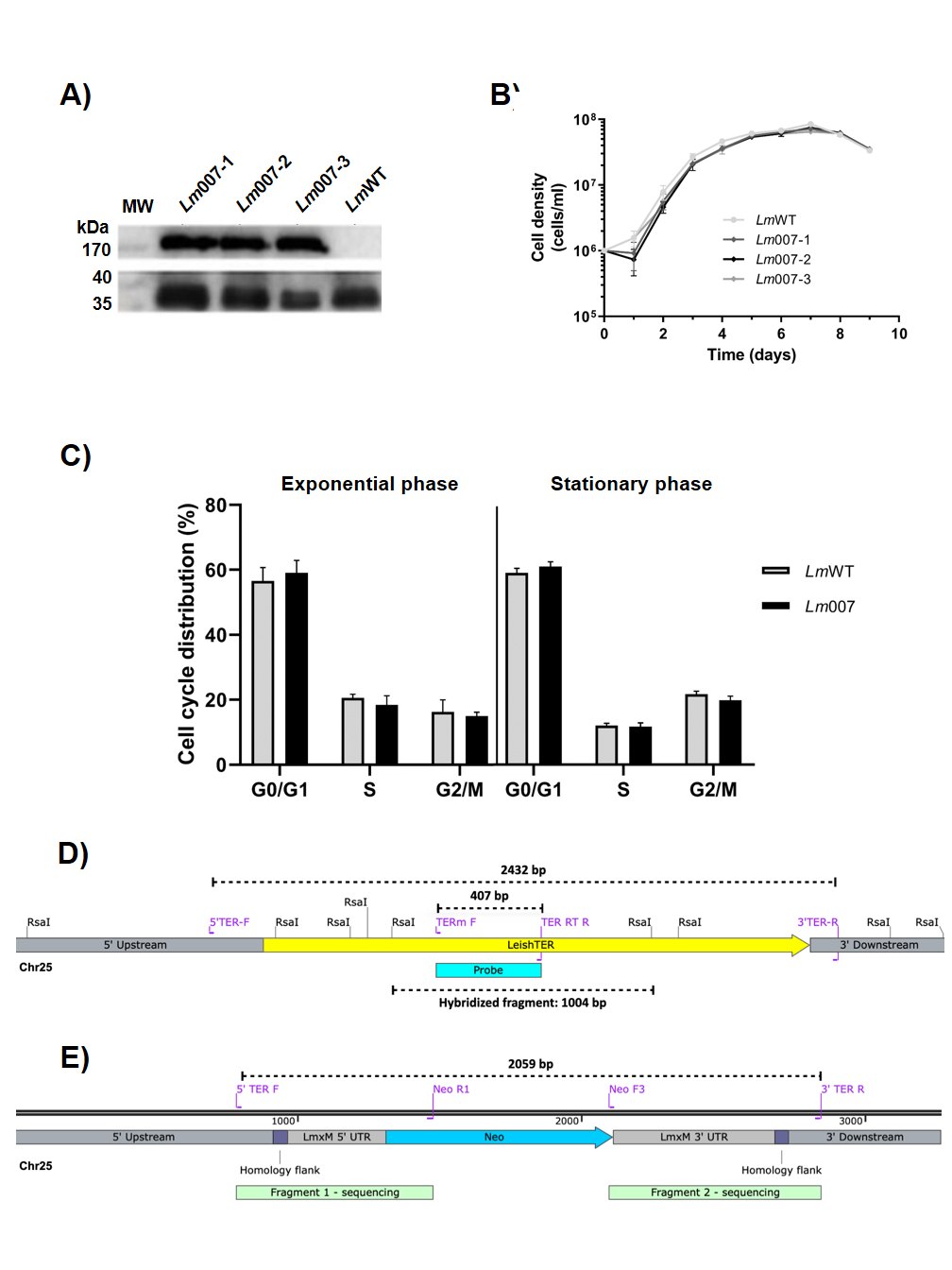

**S1 Fig. Epissomal expression of Cas9 does not interfere with proliferation and cell cycle progression in *L. major*.** A) Western blot analysis using an anti-Flag antibody was done to verify the expression of Cas9 in *Lm*007 clones. Anti-*Lm*GAPDH antibody served as the loading control, and a molecular weight marker (Page Ruler, Invitrogen) was used for reference. B) Growth curves of *Lm*007 and wild-type parasites were generated by daily cell counting (every 24 h) using a Neubauer chamber. The graph represents the mean values of technical triplicates. C) Cell cycle profiles of *Lm*007-1 clone and *Lm*WT at P5 are depicted. Cell cycle analyses were conducted on days 3 (exponential growth phase - left graph) and 5 (stationary phase - right graph). D) The schematic map illustrates the *Leish*TER locus in *Lm*WT. The position of each primer set used for PCR amplification (5' TER F + 3' TER R – S1 Table) and RT-PCR (TERm F + TER RT R – S1 Table) are signaled (purple), along with the expected amplicon sizes (2432 bp and 407 bp, respectively). The position of the *Rsa*I restriction sites, the primers used to generate the DIG-labeled TER probe (in royal blue), and the size of the fragment hybridized with the probe (1004 bp) are also shown. E) A schematic map of the *Leish*TER gene locus at chromosome 25 after gene edition using CRISPR/Cas9 is shown. The primer set (5' TER F + 3' TER R – S1 Table) was used to confirm the insertion of the donor DNA cassette that replaced *Leish*TER. The expected amplicon size (2059 bp) and the primers used for DNA sequencing analysis (5' TER F + Neo R1; Neo F3 + 3' TER R – S1 Table) are indicated.

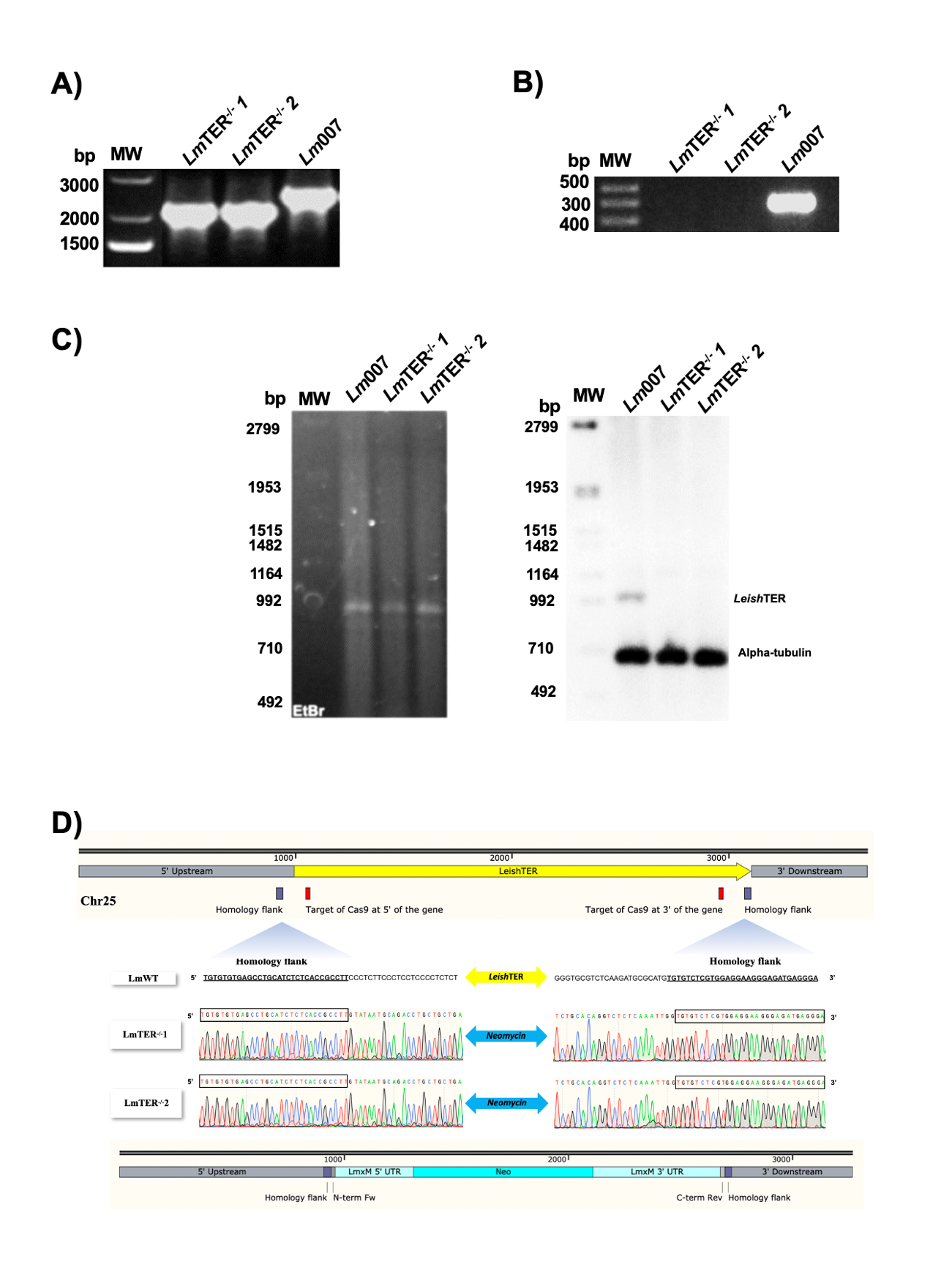

**S2 Fig. Generation of *Leish*TER double knockout (*Lm*TER^-/-^) was successfully achieved.** A) PCR amplification was performed using specific primer sets to validate the edition of the *Leish*TER locus. The PCR products were separated in a 1% agarose gel stained with ethidium bromide. A molecular weight marker (1 Kb Plus DNA ladder, Invitrogen) was included for reference. B) RT-PCR was conducted using the TERm F and TER RT R primer set (S1 Table), which amplifies an internal *Leish*TER fragment of 407 bp, and total RNA obtained from *Lm*TER^-/-^ clones and *Lm*007. The cDNA samples were fractionated in a 1% agarose gel stained with ethidium bromide. A molecular weight marker (1 Kb Plus DNA ladder, Invitrogen) was used for reference. C) Southern blot analysis was performed using *Rsa*I digested-DNA obtained from *Lm*007 and *Lm*TER^-/-^ clones, hybridized with a DIG-labeled TER probe (S1 Table). Additionally, a DIG-labeled Alpha-tubulin probe (~700 bp) served as the loading control (S1 Table). DNA molecular weight marker VII, DIG-labeled (Roche), was employed for reference. D) The diagram illustrates the outcomes of the edited and unedited *Leish*TER locus. The top image portrays the wild-type *Leish*TER locus, highlighting the homology regions (purple) and the sgRNAs target regions (red). Electropherograms (middle image) indicate the positions of the 5' and 3' homology regions containing shared sequences between *Lm*WT and *Lm*TER^-/-^ clones. The integration of the donor DNA cassette containing the neomycin resistance gene within the *Leish*TER locus (both alleles) is depicted (bottom image).

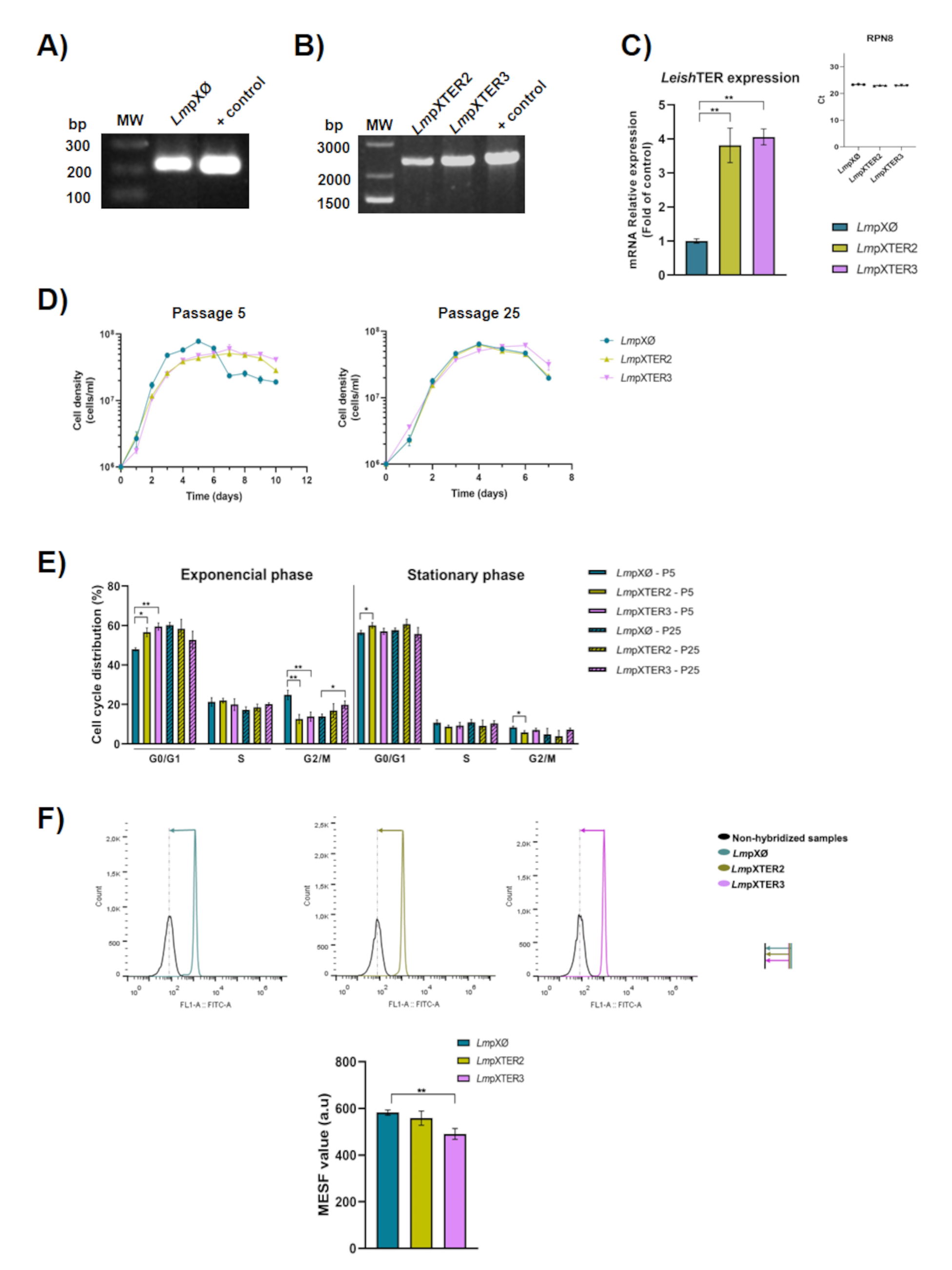

**S3 Fig. The creation of a *Leish*TER overexpressor lineage, known as *Lm*pXTER, has a dominant negative effect on the parasite similar to the depletion of *Leish*TER.** A) PCR using a specific primer set, was carried out to verify the acquisition of the control lineage carrying the pX63Neo plasmid, denoted as *Lm*pXØ. B) Clones overexpressing *Leish*TER, designated as *Lm*pXTER2 and *Lm*pXTER3, were also confirmed by PCR using the same primer set as in A). C) *Leish*TER expression was assessed by RT-qPCR using total RNA extracted from *Lm*pXTER2, *Lm*pXTER3 and *Lm*pXØ. The upper right panel displays the Ct values of the reference gene RPN8. D) Growth curves of *Lm*pXØ, *Lm*pXTER2, and *Lm*pXTER3 at P5 (left panel) and P25 (right panel) were obtained by daily counting using a Neubauer chamber. E) Cell cycle analyses of *Lm*pXØ, *Lm*pXTER2 and *Lm*pXTER3 at P5 and P25 were conducted using parasites in the exponential (left panel) and stationary growth phases (right panel). F) A PNA FITC-labeled telomeric DNA oligo probe (CCCTAA)3 (Panagene) was used to perform the Flow-FISH analysis. The histograms represent the mean fluorescence of non-hybridized parasites (in black) and parasites hybridized with the telomeric probe (*Lm*pXØ in green, *Lm*pXTER2 in yellow, and *Lm*pXTER3 in pink). Vertical lines mark the difference between peaks, and horizontal lines with arrows represent telomere length differences in the populations analyzed. The amount of fluorescence among samples was calculated using MESF, and the means ± S.D. of technical triplicate was plotted in the graph (bottom). All data are presented as mean ± SD and represent triplicate experiments. Statistical analysis was conducted using the student t-test, with Welch’s correction applied when necessary (**p* ≤ 0.05, ***p* ≤ 0.01).

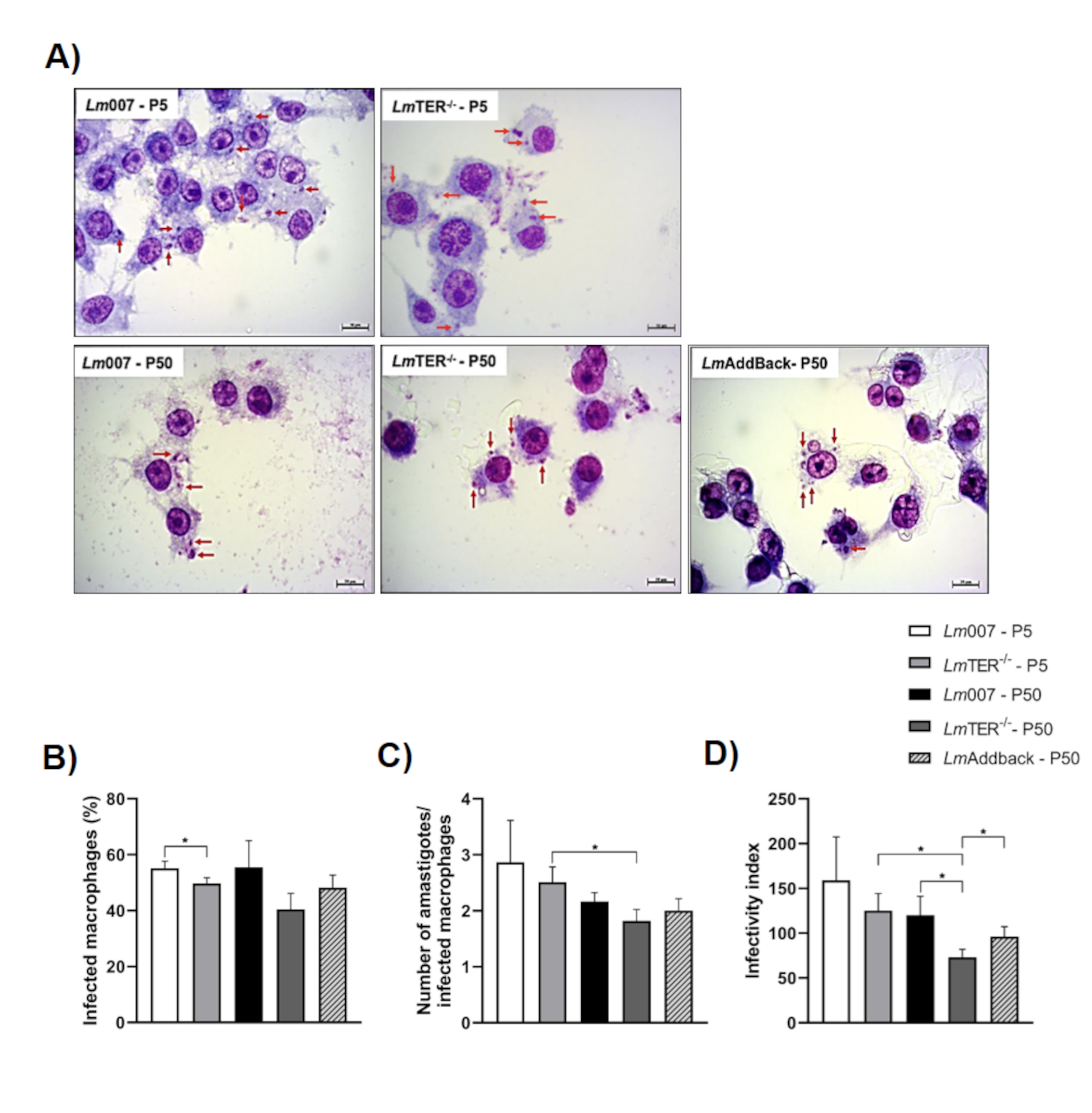

**S4 Fig. The reduced infectivity of BMDM by *Lm*TER*^-/-^* was validated through a subsequent in vitro experiment using RAW 264.7 cells.** A) Photomicrographs depict RAW 264.7 macrophages infected for 24 h with *Lm*007 and *Lm*TER^-/-^ at P5 and P50, as well as with *Lm*Addback at P50. Red arrows indicate internalized *L. major* parasites. Images were captured randomly at 1000X magnification. Scale bars = 10 µm. B) Percentage of RAW 264.7 macrophages infected with *Lm*007 and *Lm*TER^-/-^ (P5 and P50), and *Lm*AddBack (P50). C) Number of amastigotes per macrophage in infections with *Lm*007 and *Lm*TER^-/-^ (P5 and P50), and *Lm*AddBack (P50). D) The estimated infectivity index of *Lm*007, *Lm*TER^-/-^ at P5 and P50 and *Lm*AddBack at P50 in RAW 264.7 macrophages. All data are presented as mean ± SD and represent triplicate experiments. Statistical analysis was conducted using the Student t-test with Welch’s correction as needed. **p* ≤ 0.05.

**S1 Table. Oligonucleotide sequences**

| **Name** | **Sequence 5’ - 3’** | **Reference** |
| --- | --- | --- |
| For-Cas9 | CCGAAGAGGTCGTGAAGAAG | Current work |
| Rev-Cas9 | GCCTTATCCAGTTCGCTCAG | Current work |
| T7RNAP-F | TAAGCGCGGCAAGCGCCCGA | Current work |
| T7RNAP-R | GACCTACTACGCCAGCATTT | Current work |
| 5’ TER F | GGTTCTTTCCCCTTGCACTG | Current work |
| 5’ TER R | CAATCTCCCGTGTGGTGCTT | Current work |
| 3’ TER R | TTCTGCGGTTCAGCACTACA | Current work |
| Neo R1 | AGGTCGGTCTTGACAAAAAGAACC | Current work |
| Neo F3 | TGACGAGTTCTTCAGCTCCG | Current work |
| TracrRNA (G00) – scaffold | AAAAGCACCGACTCGGTGCCACTTTTTCAAGTTGATAACGGACTAGCCTTATTTTAACTTGCTATTTCTAGCTCTAAAAC | [34] |
| TER-seed 5’ | GAAATTAATACGACTCACTATAGGAGGTGTCGGACGATGTCCTAGTTTTAGAGCTAGAAATAGC | Current work, generated at http://grna.ctegd.uga.edu |
| TER-seed 3’ | GAAATTAATACGACTCACTATAGGAGCGTGGCGTCGATAAAAACGTTTTAGAGCTAGAAATAGC | Current work, generated at http://grna.ctegd.uga.edu |
| N-term For (TER) | TGTGTGTGAGCCTGCATCTCTCACCGCCTTGTATAATGCAGACCTGCTGC | Current work, generate at http://www.leishgedit.net/Home.html |
| C-term Rev (TER) | TCCCTCATCTCCCTTCCTCCACGAGACACACCAATTTGAGAGACCTGTGC | Current work, generate at http://www.leishgedit.net/Home.html |
| DIG-labeled TelC probe | CCCTAACCCTAACCCTAACCCTAACCCTAA | [41, 81] |
| Lmj 18s RNA Fw | AGTGTGGAGATCGAAGATGATTAG | Current work |
| Lmj 18s RNA Rv | CCCTGAGACTGTAACCTCAAAG | Current work |
| DIG-labeled 18S rRNA probe | AGTGTGGAGATCGAAGATGATTAGAGACCATTGTAGTCCACACTGCAAACGATGACACCCATGAATTGGGGATCTTATGGGCCGGCCTGCGGCAGGGTTTACCCTGTGTCAGCACCGCGCCCGCTTTTACCAACTTACGTATCTTTTCTATTCGGCCTTTACCGGCCACCCACGGGAATATCCTCAGCACGTTTTCTGTTTTTTCACGCGAAAGCTTTGAGGTTACAGTCTCAGGG | Current work |
| AlfaTub-1 Fw | CGAACAACTACGCTCGTGGC | Current work |
| AlfaTub-1 Rv | GCCACCCACAGCGTGGAACA | Current work |
| DIG-labeled Alpha-Tubuline probe | CGAACAACTACGCTCGTGGCCACTACACGATCGGCAAGGAGATCGTCGACCTTGCGCTGGACCGCATTCGCAAGCTGGCGGACAACTGCACTGGTCTCCAGGGCTTTATGGTGTTCCACGCTGTGGGTGGC | Current work |
| TERm F | TGCTTGCATGTCTGTCCTTC | Current work |
| TER RT F | GAGAACTAGCACGGCCACTC | Current work |
| TER RT R | GAGGATCGCGCTACAAAGTC | Current work |
| DIG-labeled TER probe | TGCTTGCATGTCTGTCCTTCGTCTCAGCACGCGCGTGCGCGCACACTATGCCGGCCCTTCATCGCTCAACGGCTCTTTTTCCGTTTCGCACCTGTGCCCTCAGGAGCGCGTAAGCACGCCTAAGCACACACCGTCACGCAGAATCAGACCCCACGCGTGATTGCAAGCGATGCGGCCTCTCACTGCAGTCGAGGTGGGGCAACGGGCGCAGTCCGCCTTCTCCCTTTCTCATTTGCTTTTCAGTATGCTGGCCTTCTCGCCGTCGGTGAGGGTGTTATGTGTGTGCGTGCGGGCGTGTGGTGCACCACCATGGACTGTGAGAACTAGCACGGCCACTCAGGCATGCACACATACACGTGCACGTGCACGTGCACACGCGCGTGCACCGACTTTGTAGCGCGATCCTC | Current work |
| SUBCHRO. 29R-F | ATGGGGATTAAGGGAAGCAC | [38] |
| LMCHR29_RV_2 | CCCGTACCCGTACCTTATCC | Current work |
| RPN8 F | ATGAACCGCCGCAAGCT | [38] |
| RPN8 R | GGCGCGACGACGATCTTTGATT | [38] |
| pX63 Neo Fw | CACCACCCTCAACCACCC | Current work |
| pX63 Neo Rv | CAATACGCAAACCGCCTC | Current work |

**S2 Table. Flow-FISH (MESF ± SD analysis) of *Lm*007, *Lm*TER^-/-^, *Lm*pXØ and *Lm*pXTER cells**

| **Samples** | **MESF value (a.u.) ± SD** | **Proportional change (MESF analysis)** |
| --- | --- | --- |
| *Lm*007 - P5 | 516.3 ± 34.65 | 1 |
| *Lm*TER^-/-^ - P5 | 354.3 ± 13.43 | 0,69 |
| *Lm*007 - P50 | 502.7 ± 30.83 | 1 |
| *Lm*TER^-/-^ - P50 | 261.7 ± 14.29 | 0,52 |
| *Lm*pX0 - P25 | 582.3 ± 11.02 | 1 |
| *Lm*pXT2 - P25 | 558.7 ± 30.57 | 0,96 |
| *Lm*pXT3 - P25 | 491 ± 23.07 | 0,84 |

**S3 Table. In Vitro Infectivity Assessment of *Lm*TER^-/-^, *Lm*007 (P5 and P50), and *Lm*AddBack (P50) on Balb/c Bone Marrow-Derived Macrophages (BMDM)**

| **Samples** | **Infected macrophages (%) ± S.D.** | **Number of amastigotes/infected macrophages ± S.D.** | **Infectivity index (%) ± S.D.** |
| --- | --- | --- | --- |
| *Lm*007 – P5 – 24 h | 29.8 ± 1.96 | 1.86 ± 0.17 | 55.4 ± 6.67 |
| *Lm*TER^-/-^ - P5 – 24 h | 18.5 ± 0.61 | 1.65 ± 0.37 | 30.7 ± 7.65 |
| *Lm*007 – P50 – 24 h | 31.5 ± 2.27 | 2.4 ± 0.22 | 75.7 ± 9.62 |
| *Lm*TER^-/-^ - P50 – 24 h | 27.9 ± 4.05 | 1.81 ± 0.04 | 50.5 ± 6.21 |
| *Lm*Addback – P50 – 24 h | 36.1 ± 3.49 | 1.74 ± 0.16 | 63 ± 8.09 |
| *Lm*007 – P5 – 48 h | 69.2 ± 6.52 | 3.35 ± 0.24 | 232 ± 28.7 |
| *Lm*TER^-/-^ - P5 – 48 h | 28.5 ± 1.95 | 1.91 ± 0.08 | 54.3 ± 1.48 |
| *Lm*007 – P50 – 48 h | 57.8 ± 1.02 | 3.18 ± 0.20 | 184 ± 12.1 |
| *Lm*TER^-/-^ - P50 – 48 h | 50.8 ± 3.42 | 2 ± 0.03 | 102 ± 6.42 |
| *Lm*Addback – P50 – 48 h | 44.2 ± 4.19 | 2.11 ± 0.14 | 92.9 ± 6.71 |
